# Associations Between Gut Microbiome and 24-Hour Blood Pressure Variability: A Cross-sectional Study Highlighting Sex Differences and Potential Therapeutic Targets

**DOI:** 10.1101/2025.11.03.685706

**Authors:** Preeti Dinesh Virwani, Gordon Qian, Carman Nga-Man Cheung, Tommy K.K.T.S. Pijarnvanit, Matthew S.S. Hsu, Yick Hin Chow, Lok Kan Tang, Yiu-Hei Tse, Jia-Wen Xian, Shirley Sau-Wing Lam, Crystal P.I. Lee, Chelsea C.W. Lo, Roxanna K.C. Liu, Tsi Lok Ho, Bak Yue Chow, Kin Sum Leung, Emily K.K. Lo, Man-Fung Yuen, Suet Yi Leung, Ivan Fan-Ngai Hung, Jimmy Chun Yu Louie, Kay-Cheong Teo, Hani El-Nezami, Joshua Wing Kei Ho, Kui Kai Lau

## Abstract

Blood pressure (BP) variability is an independent risk factor for cardiovascular disease (CVD). While gut microbiota (GM) and GM-derived short-chain fatty acids (SCFAs) are recognized to play a key role in BP regulation, their association with BP variability remains poorly understood. This cross-sectional study of 241 community-dwelling individuals from Hong Kong (113 men and 128 women, mean age 54±6 years) without symptomatic CVD examined the associations of GM, characterized by shotgun sequencing, and plasma SCFAs, with systolic and diastolic blood pressure (SBP/DBP) variability assessed by 24-hour BP monitoring. GM analyses, using covariate-adjusted statistical models, revealed that higher 24-hour SBP CoV associated negatively with GM α-diversity (Shannon and Simpson’s index, *P*<0.05) and positively with Firmicutes/Bacteroidetes ratio, driven by the female cohort. *Parabacteroides merdae*, *Bacteroides dorei*, *Bifidobacterium pseudocatenulatum*, *Alistipes finegoldii* and *Bacteroides intestinalis* were negatively associated with indices of SBP/DBP variability in a sex-specific manner. Further analysis indicated that *B. dorei* may mediate SBP CoV via plasma iso-butyric acid in women (bootstrapping 95% CI: −3.6 to −0.19; *P*<0.05). We demonstrated that higher SBP variability is associated with markers of gut dysbiosis and a reduction in beneficial gut bacteria, particularly in women. Notably, we identified several gut bacterial species with potential therapeutic implications for managing 24-hour SBP/DBP variability, warranting further investigation.

## Introduction

Cardiovascular diseases (CVDs) are the leading cause of morbidity and mortality taking approximately 18 million lives a year worldwide^1^. While high blood pressure (BP) is the leading risk factor for CVD, BP variability has also been associated with worse CVD outcomes and mortality independent of mean BP^1,2^. Disruption of 24-hour BP fluctuation rhythm is linked to endothelial dysfunction and cardiovascular organ damage, with clinical analyses indicating that BP variability predicts cardiovascular events, adverse stroke outcomes, and all-cause mortality^2–5^. Moreover, the beneficial effects of certain classes of antihypertensive agents in reducing secondary stroke risk may be partly attributed to their BP variability lowering effects^6^. Taken together, increased BP variability is an independent modifiable risk factor for cardiovascular diseases and potentially a therapeutic target.

The diverse array of microorganisms inhabiting the gastrointestinal tract, collectively referred to as gut microbiota (GM), plays a crucial role in influencing cardiovascular health^7^. GM is recognized as a key modulator of BP, supported by compelling evidence from both animal and human studies^8–10^. GM may mediate BP through its metabolites, short-chain fatty acids (SCFAs) which bind to receptors located on kidneys, blood vessels and immune cells; SCFAs regulate BP via multiple mechanisms promoting vasodilation and reducing systemic inflammation^9^. Recent studies indicate that GM-based therapies, such as probiotics and SCFA supplements, effectively reduce BP in humans, highlighting their therapeutic potential in BP regulation^9,10^. However, despite a plethora of investigations into the associations between GM and BP, it remains unclear if GM plays a role in modulating BP variability. The composition of gut microbiome exhibits rhythmic fluctuations over a 24-hour period that align with rhythmic variations in BP within the same timeframe^11^. This correlation suggests that GM may mediate 24-hour BP variability. To date, there is only one clinical study that has reported a plausible association between GM and systolic BP variability in a mixed-sex cohort^12^. This study utilized 16S rRNA gene sequencing, which is known for its low resolution in species annotation, and did not adjust for confounding factors, particularly mean BP and sleep quality, which can significantly influence GM^12,13^. The disruption of sleep patterns and host molecular clock proteins have been associated with GM dysbiosis^14^, underscoring the importance of adjusting for sleep quality parameters in studies investigating the role of GM in pathophysiology of abnormal BP variability.

Whilst we have previously shown sex differences in association between GM and hypertension^15^, whether there are sex-specific associations in GM and BP variability remains to be investigated. Moreover, it is still unknown if GM is associated with diastolic BP variability, which has been reported as an independent predictor of cardiovascular organ damage and worse CVD outcomes^5^. To address these gaps, we conducted a cross-sectional study to investigate the link between GM profiled by shotgun metagenomic sequencing and 24-hour BP variability assessed by 24-hour SBP/ DBP CoV, and nighttime dipping status. Sex-stratified analysis was conducted under different statistical models adjusting for various confounding factors^15–20^.

## Methods

### Study design and population

This study was approved by the Institutional Review Board of the University of Hong Kong/ Hospital Authority Hong Kong West Cluster (UW 18-498) and informed consent was obtained from all participants. 284 individuals residing in Hong Kong were enrolled between July 2018 and June 2020 through community recruitment by open advertisements. The study design was as previously detailed^15^. Briefly, inclusion criteria were age 40 to 65 years, Chinese ethnicity, with no known symptomatic cardiovascular, neurological, gastrointestinal, neurodegenerative, autoimmune diseases or malignancy. Participants with recent gastroenteritis, febrile illness, steroids, antibiotics or probiotics use within 4 weeks and recent travel to tropical areas within 6 months were excluded. After excluding 43 participants who took anti-hypertensive agents, 241 participants were included in the final analysis.

### 24-hour BP measurement and calculation of BP variability

A calibrated ambulatory BP monitoring device, TM-2441 (A&D Medical, Japan) was fitted onto the participants, with the BP cuff placed on their non-dominant arm to record the 24-hour ambulatory BP. The BP was measured every 30 minutes from 6 am to 10 pm and then every 60 minutes from 10 pm to 6 am. Variability in BP was assessed by estimating SBP and DBP coefficient of variation (CoV) and nighttime dipping status. Participants were asked to record the wake-up and sleep time to determine the nighttime systolic BP dipping status. The sleep time BP was calculated by averaging the BP readings from the time the participant slept until he/she woke up and the awake BP was the average of BP readings during the awake time^21^. Participants were classified according to the percentage of dip in their nighttime SBP [100×(1−sleep SBP/awake SBP)] as extreme-dippers (≥20%), dippers (≥10% but <20%), nondippers (≥0% but <10%) and reverse-dippers (<0%)^21^.

### Stool collection, DNA extraction, library preparation and sequencing

Each participant was given a stool collection kit (2-piece sample collector, DYND36500, Medline Industries Inc., USA) for stool collection at home. The stool collected in the sample container was refrigerated and returned to the laboratory within 24 hours in the provided ice bag and ice packs. The stool samples were homogenized, subsampled into 4 equal aliquots and stored at −80°C until further processing. DNA extraction from the stool and shotgun sequencing methods were as previously described^15^.

### Taxonomic and functional profiling of raw sequencing data

Sequencing read pre-processing and analysis methods were as previously detailed^15^. Briefly, raw sequencing reads were quality filtered and trimmed using Bbduk from the BBmap Suite^22^. Contaminant sequences were removed by aligning to human and PhiX reference genomes. Species abundances were estimated using MetaPhlAn 3.0^23^ and gene family abundances were estimated using HUMAnN3 and then regrouped into Kyoto Encyclopedia of Genes and Genomes (KEGG) ontology higher-level functional annotations^24^.

### Quantification of plasma SCFAs

The SCFAs, acetic acid, propionic acid, butyric acid, and iso-butyric acid were quantified using Agilent 7890B gas chromatography-mass spectrometry (Agilent Technologies, Santa Clara, CA USA) as previously described^25,26^.

### Assessments of covariates and dietary intake

Fasting blood samples were drawn from the participants. Serum total cholesterol, triglycerides, HDL and LDL levels were estimated using commercial kits (TECO Diagnostics, USA #C507-480, #T532-1L, #H513-100, #L530-100). Serum glucose was measured using a glucose meter, ACCU-CHEK Performa (Roche, USA). To estimate sleep parameters, participants wore an Actigraphy watch (Actigraph, LA) for 7 days and were instructed to only take off the watch when they took a shower and when they swam. Sleep and wake-up times were estimated by the Actigraph watch, and sleep latency and sleep efficiency were automatically estimated using the Actigraph software. The presence of hepatic steatosis was assessed using a fibroscan (EchosensTM, Paris, France). The controlled attenuation parameter (CAP) score was used to measure the severity of steatosis. Cutoffs for mild, moderate, and severe steatosis were 248-267, 268-279 and ≥280 dB/M respectively^27^. The urinary sodium and urinary creatinine were measured in the second morning spot urine using commercial kits (Abcam, UK #ab 211096, #ab 204537) to estimate the daily sodium intake as described by Kawasaki et al^28^. The methods for dietary intake assessment based on weekly food recall were described previously^15^.

### Statistical and Bioinformatics analysis

Statistical analyses were performed using the R statistical environment (version 4.3). The BP variability data analyses were carried out using independent sample *t*-tests or *χ*^2^ tests as appropriate and presented data as mean ± SD for continuous variables and absolute numbers with percentages for categorical variables. The SCFA data were analyzed using GraphPad Prism (version 9.2.0) and outliers detected by default Robust regression and outlier removal (ROUT) method were removed from the analysis. The correlation was determined by Pearson or Spearman depending on the distribution of the data. A two-sided *P* value < 0.05 was considered statistically significant.

Taxonomic groups were filtered based on a minimum relative abundance threshold of more than 0.1% abundance at a prevalence of 10%. Relative abundances were renormalised for each sample to sum to one. Further arcsine square-root normalisation was applied for all downstream statistical abundance analyses. *α-* and *β*-diversity indexes, Bray Curtis distance matrix, and Permutational Multivariate Analysis of Variance (PERMANOVA) analysis were performed using the *vegan* package^29^. Principle coordinates analysis (PCoA) was performed using classical multidimensional scaling (MDS) with Bray Curtis distances. Statistical association testing was performed using logistic and linear regression models which allowed adjustment of confounding variables^15–20^. In differential abundance testing of GM species, a stricter P value of less than 0.01 was considered significant. The following covariate adjustment models (m) were applied to these statistical tests: m1: no covariate adjustment; m2: age, sex and BMI; m3: m2 + sodium intake estimated by using spot urine analysis, smoking and menopause status, blood glucose, triglycerides, LDL and HDL cholesterol, hepatic steatosis via CAP score obtained by liver fibroscan; m4: m3 + sleep latency and mean 24-hour SBP or DBP. Mediation analysis was applied to test whether gut microbial associations with blood variability measurements were mediated by other variables. This was performed using the mediation library in R, where paths were assessed using linear regression with the inclusion of covariates of model 4. Significance of unstandardized indirect effects was computed through bootstrapping procedures (n=1000), and the 95% confidence interval was computed by determining the indirect effects at the 2.5th and 97.5th percentiles.

## Results

### Baseline clinical characteristics and sex differences in BP variability

The baseline clinical characteristics of 241 participants (mean age 54 ± 6 years; 53.1% females) and comparisons between males and females are described in Table 1. Women had a significantly higher 24-hour SBP CoV [mean (SD), 11(2.9) vs. 9.9(1.8); *P*<0.001] and DBP CoV [mean (SD), 14.7(3.9) vs. 13.2(3.2); *P*<0.01] than men, driven by the daytime CoV (all *P*<0.001; Figs. 1a). The difference in 24-hour SBP/DBP CoV in women vs. men remained strongly significant after adjusting for various confounding factors in our covariate-adjusted models (all *P*<0.05-0.001 in m1-m4; Table S1). Analysis of nocturnal dipping status showed that nondippers were most prevalent in our cohort (54%), though no significant sex differences were detected in the dipping status between men and women (Table 1). Dietary data analyses showed that none of the macronutrients, including fiber, carbohydrates, fat, and protein were significantly associated with BP variability indices in the whole cohort or sex-stratified analysis (data not shown; all p>0.05).

**Fig. 1.**
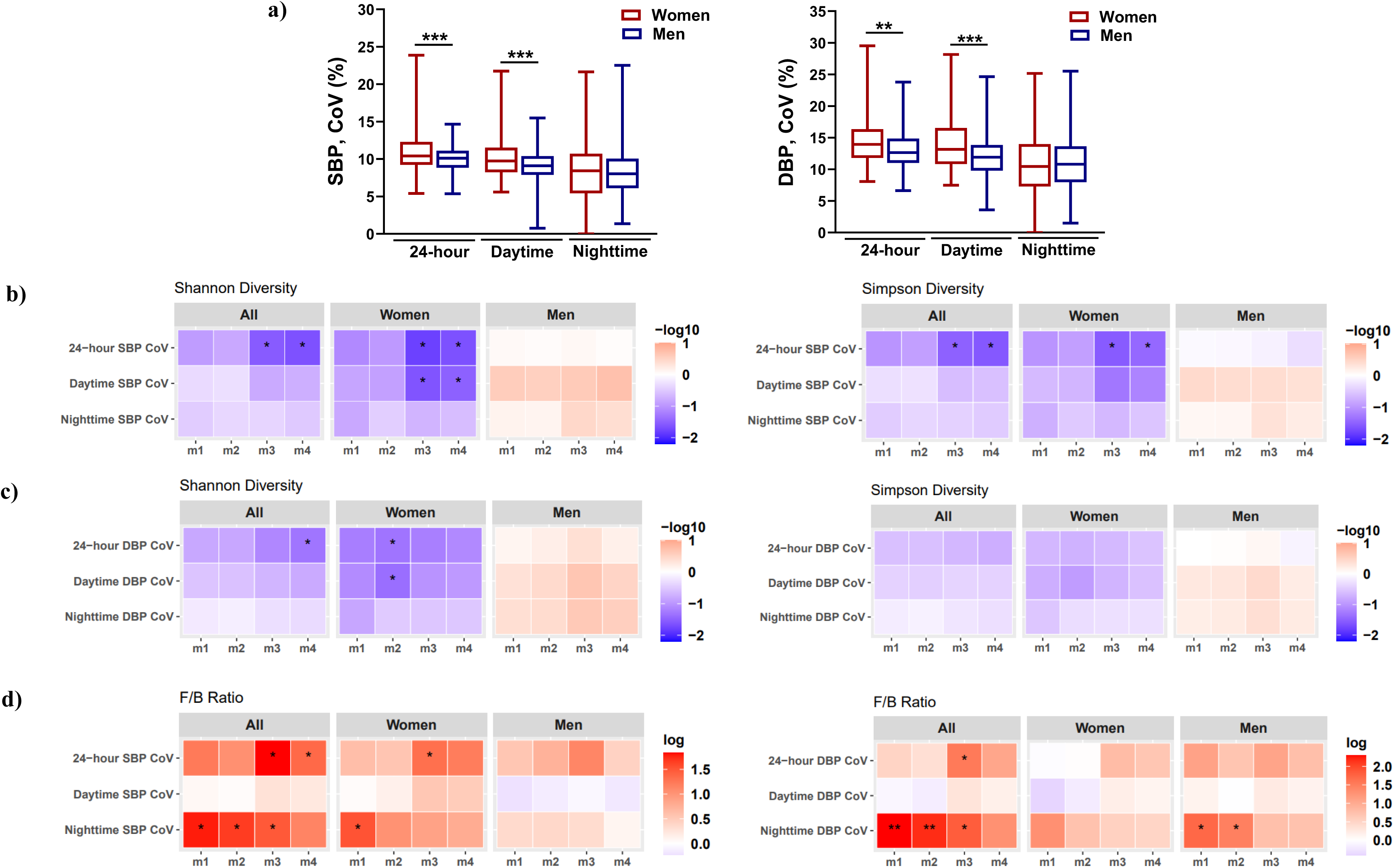
Sex differences in 24-hour, daytime, and nighttime BP CoV and markers of GM dysbiosis. **(a)** Box plots showing 24-hour, daytime and nighttime SBP (left panel)/ DBP (right panel) CoV in women vs. men. The statistical significance was determined by unpaired t-test: \*\**P* < 0.01, \*\*\**P* < 0.001. **(b-c)** Association between Shannon diversity (left panel) and Simpson’s diversity (right panel) with 24-hour, daytime and nighttime **(b)** SBP and **(c)** DBP CoV. **(d)** Association of F/B ratio with 24-hour, daytime and nighttime SBP (left panel) and DBP (right panel) CoV. All data (b-d) were analyzed by linear regression analysis after adjusting for confounding factors under the four models (m) of covariate adjustment, m1: no covariate adjustment, m2: age; sex; BMI, m3: m2 + sodium intake based on spot urine analysis; serum glucose, triglyceride, HDL and LDL cholesterol; smoking and menopause status; liver steatosis by CAP score, m4: m3 +. 24-hour mean SBP or DBP and sleep latency. The statistically significant *p* values are indicated as \**P* < 0.05, \*\**P* < 0.01.

**Table 1.** Clinical characteristics of the study population. Legend. Hypertensive status was based on 24-hour ambulatory BP. The data are represented as mean ± standard deviation. Statistical significance was assessed by unpaired t test; *P<0.05, †P<0.01, ‡P<0.001, §P<0.0001, vs. normotensive within all cohort, male and female groups. BMI, body mass index; CAP, controlled attenuation parameter; DBP, diastolic blood pressure; SBP, systolic blood pressure; WHR, waist-to-hip ratio; HDL, high-density lipoprotein; LDL, low-density lipoprotein.

| Variables | All (n=241) | Males (n=113) | Females (n=128) |
| --- | --- | --- | --- |
| Age | 54 ± 6 | 55 ± 6 | 54 ± 6 |
| BMI | 23.4 ± 3.3 | 24.6 ± 3.1 | 22.3 ± 3.1§ |
| WHR | 0.86 ± 0.07 | 0.89 ± 0.05 | 0.82 ± 0.07§ |
| Clinic SBP, mmHg | 120 ± 15 | 125 ± 15 | 115 ± 14§ |
| Clinic DBP, mmHg | 77 ± 10 | 81 ± 9 | 73 ± 9§ |
| 24-hour SBP, mmHg | 122 ± 13 | 127 ± 13 | 117 ± 12§ |
| 24-hour DBP, mmHg | 76 ± 10 | 81 ± 9 | 71 ± 8§ |
| Daytime SBP, mmHg | 124 ± 13) | 129 ± 13 | 120 ± 12§ |
| Daytime DBP, mmHg | 77 ± 10 | 83 ± 10 | 73 ± 9§ |
| Nighttime SBP, mmHg | 115 ± 16 | 121 ± 14 | 109 ± 16§ |
| Nighttime DBP, mmHg | 70 ± 11 | 76 ± 10 | 65 ± 10§ |
| Hypertension staging by 24-hour BP, n (%) |  |  |  |
| Normotensive | 154 (64) | 51 (45) | 103 (80) § |
| Hypertensive | 87 (36) | 62 (55) | 25 (20) § |
| White coat hypertension | 5 (2) | 1 (1) | 4 (3) |
| Masked hypertension | 54 (22) | 35 (31) | 19 (15) † |
| Dipping status, n (%) |  |  |  |
| Extreme dippers | 7 (3) | 1 (1) | 6 (5) |
| Dippers | 70 (29) | 28 (25) | 42 (33) |
| Nondippers | 129 (54) | 66 (58) | 63 (50) |
| Reverse dippers | 34 (14) | 18 (16) | 16 (13) |
| Smoking status, n (%) |  |  |  |
| Current smokers | 12 (5.0) | 12 (10.6) | 0 |
| Ex-smokers | 37 (15.4) | 29 (25.7) | 8 (6.3) |
| Sodium intake estimated by spot urine analysis, g/d | 5.48 ± 1.68 | 5.69 ± 1.66 | 5.30 ± 1.68 |
| Total cholesterol, mmol/L | 5.08 ± 0.88 | 4.98 ± 0.77 | 5.17 ± 0.96 |
| HDL-cholesterol, mmol/L | 1.47± 0.26 | 1.37 ± 0.23 | 1.56 ± 0.26 § |
| LDL-cholesterol, mmol/L | 2.58 ± 0.52 | 2.53 ± 0.51 | 2.62 ± 0.52 |
| Triglyceride, mmol/L | 1.06 ± 0.71 | 1.20 ± 0.85 | 0.94 ± 0.54† |
| Fasting glucose, mmol/L | 5.7 ± 0.9 | 5.8 ± 1.0 | 5.6 ± 0.7 |
| Sleep latency, min | 7.3 ± 5.9 | 7.4 ± 6.3 | 7.3 ± 5.6 |

### Gut microbiota composition and 24-hour SBP and DBP variability

Whole-genome shotgun sequencing of 241 stool samples yielded 38.68 ± 9.97 (mean ± SD) million reads per sample after filtering. The GM beta(β)-diversity (between-person) was not significantly associated with SBP/DBP CoV (data not shown). There was a negative association between microbial Shannon and Simpson’s diversity index and 24-hour SBP CoV in the whole cohort, which was significant after adjusting for confounding variables in m3 (age, sex, BMI, serum glucose and lipids, smoking and menopause status, fatty liver CAP score, sodium intake estimated by urine analysis) and remained significant after further adjusting for sleep latency and mean SBP in m4 (all *P*<0.05; Fig. 1b, Table S2). Sex-stratified analyses indicated that the associations between *α*-diversity measures were driven by the female cohort (all *P*<0.05 in m3-m4; Fig. 1b, Table S2), suggesting reduced GM α-diversity in women with greater systolic BP variability. However, DBP CoV was not significantly associated with the measures of α-diversity except significant negative association of Shannon index with 24-hour DBP CoV in the whole cohort only after additional adjustment for mean SBP and sleep latency under m4 (*P*<0.05; Fig. 1c; Table S2). Similarly, Firmicutes/Bacteroidetes (F/B) ratio had a significant positive association with the 24-hour SBP CoV, but not DBP CoV, after adjusting for confounding variables under m3-4 in the whole cohort (all *P*<0.05; Fig. 1d; Table S3). Taken together, these results suggested that participants with higher 24-hour systolic BP variability may have gut dysbiosis.

Shotgun sequencing analysis identified several bacterial species with significant sex-specific associations with 24-hour, daytime, and nighttime systolic and diastolic BP variability indices (Table S4). *Parabacteroides merdae* had a significant negative association with 24-hour SBP/DBP CoV in the whole cohort, which remained upon adjustments with various confounding factors in all of the covariate-adjusted models (all *P*<0.01; Fig. 2a, left panel; Table S4). Sex-stratified analysis showed that *P. merdae* was significantly reduced in both men and women with higher systolic BP CoV (*P*<0.05-0.01; Fig. 2a, left panel; Table S4). All of the other significant associations between gut bacterial species and various indices of systolic and diastolic BP variability were sex-specific (Fig. 2a; Table S4). *Bacteroides dorei* was significantly reduced in women with higher 24-hour SBP CoV, driven by daytime SBP (*P*<0.05-0.01 in m1-m4; Fig. 2a, left panel; Table S4). Other significant associations with systolic BP variability in women included negative association of *Paraprevotella xylaniphila* with daytime SBP CoV, and *Coprococcus catus*, *Blautia wexlerae*, and *Bacteroides plebius* with nighttime SBP CoV (*P*<0.05-0.01 in m1-m4; Table S4). In men, *Bifidobacterium pseudocatenulatum* had a significant negative association with 24-hour SBP CoV (*P*<0.05-0.01 in m1-m2, m4; Figs. 2a, left panel; Table S4), whereas *Ruminococcus bicirculans* and *Roseburia intestinalis* were enriched in men with higher nighttime SBP CoV (*P*<0.05-0.01 in m1-m4; Table S4).

**Fig. 2.**
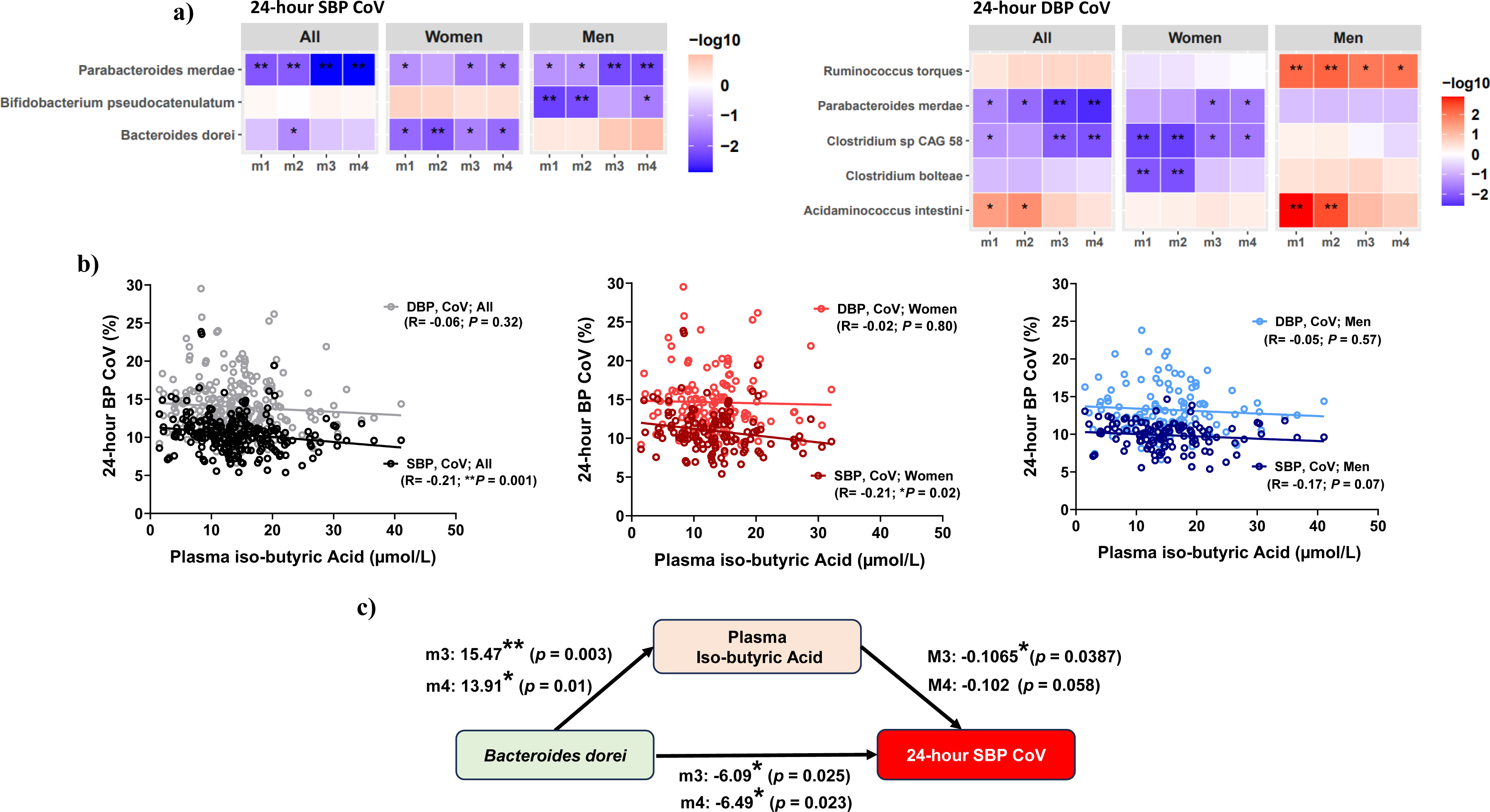
Gut microbiota species and short-chain fatty acids associated with 24-hour SBP and DBP CoV in the whole cohort and sex-stratified analyses. Gut bacterial species associated with **(a)** 24-hour SBP (left panel) and DBP (right panel) CoV through linear regression analysis under 4 different models for covariate adjustment. Only species with at least < 0.01 significance in at least one model are shown. **(b)** Scatterplot correlation between 24-hour SBP/DBP CoV and plasma iso-butyric acid in the whole cohort (left panel), women (middle panel) and men (right panel). **(c)** Mediation of 24-hour SBP CoV by *Bacteroidetes dorei* via plasma iso-butyric acid in women. Casual mediation analysis was carried out using nonparametric bootstrap confidence intervals with the percentile method and 1000 simulations. Covariate-adjusted models (m), m1: no covariate adjustment, m2: age; sex; BMI, m3: m2 + sodium intake based on spot urine analysis; serum glucose, triglyceride, HDL and LDL cholesterol; smoking and menopause status; liver steatosis by CAP score, m4: m3 +. 24-hour mean SBP or DBP and sleep latency. The statistically significant *p* values are indicated as \**P* < 0.05, \*\**P* < 0.01.

Significant GM associations with diastolic BP variability included reduction of *P. merdae* and *Clostridium* sp CAG 58 with higher 24-hour DBP CoV in the whole cohort, driven by women (all *P*<0.05-0.01; Fig. 2a, right panel; Table S4). Furthermore, *Roseburia intestinalis* had a strongly significant negative association with daytime DBP CoV (*P*<0.01-0.001 in m1-m4; Table S4) and *Fusicatenibacter saccharivorans* with nighttime DBP CoV in women (all *P*<0.0 in m2-m4; Table S4). The GM of men with higher DBP CoV was significantly enriched in *Ruminococus torques* (*P*<0.05-0.01 in m1-m4; Fig. 2a, right panel; Table S4).

### GM-derived short-chain fatty acids and 24-hour SBP and DBP variability

GM metabolites, SCFAs have been recognized for a role in BP regulation via their receptors located on kidneys, blood vessels and immune cells^9,10^. Therefore, we investigated if plasma levels of SCFAs were associated with indices of 24-hour BP variability. Plasma iso-butyric acid had a significant negative correlation with 24-hour SBP CoV in the whole cohort (R = −0.21, *P*=0.001; Fig. 2b, left panel). Upon stratification by sex, plasma iso-butyric acid was significantly correlated with 24-hour SBP CoV in women (R = −0.21; *P*=0.02; Fig. 2b, middle panel), but not in men (R = - 0.17; *P*=0.07; Fig. 2b, right panel). The significance of the associations between plasma SCFAs and BP variability indices persisted after adjusting for various confounding factors in the statistical models of covariate adjustments (Table S5).

We further investigated the relationship between plasma SCFAs and GM species that were significantly associated with BP variability indices in our cohort (Table S6). Notably, *B. dorei* had a significant positive correlation with plasma iso-butyric acid in women (*P*<0.05-0.01 in m1-m4; Table S6). A statistical mediation analysis showed that the effect of *B. dorei* on 24hr SBP co-variability was mediated via plasma iso-butyric acid after adjusting for confounding factors. As Figure 2d illustrates, the regression coefficient between *B. dorei* and 24hr SBP co-variability and the regression coefficient between plasma iso-butyric acid and 24hr SBP co-variability was significant. The indirect effect was (15.47)*(−0.1065) = 1.64 in m3 and (13.91)*(−0.102) = −1.41 in m4 after further adjusting for mean 24-hour SBP and sleep latency. The bootstrapped unstandardized indirect effect was −1.64, and the 95% confidence interval ranged from −3.6 to −0.19 in m3 and −1.41, and the 95% confidence interval ranged from −3.49 to −0.03 in m4. Thus, the indirect effect was statistically significant (p<0.05) (Fig. 2d; Table2).

### Gut microbiome/SCFAs and nighttime dipping

The GM beta diversity did not significantly differ between participants classified as dippers compared to nondippers, extreme and reverse dippers (data not shown). However, there was a significant positive association between Shannon and Simposon’s index and nondipping status in the whole cohort, driven by men (all *P*<0.05-0.01 in m1-m4; Figs. 3a). *Bacteroides intestinalis* was significantly reduced in nondippers compared to dippers in the female cohort, which remained after adjusting for various confounding factors in m2-m4 (all *P*<0.01; Fig. 3b; Table S7). Additionally, sex-stratified analyses revealed that women and men with nondipping status had reduced *Alistipes finegoldii* and increased abundance of *Eubacterium rectale*, respectively, which became significant after adjusting for covariates in m2-m4 (*P*<0.05-0.01; Fig. 3b; Table S7). While *Butyricimona virosa* was significantly reduced in the extreme dippers compared to dippers, nondippers and reverse dippers in the whole cohort, driven by women (all *P*<0.05; Fig S1), this association did not remain significant after adjusting for confounding factors. Analysis of SCFAs showed that plasma butyric acid levels were significantly lower in men with normal dipping status than nondippers (*P*<0.05; Fig. 3c), which remained significant in all covariate-adjusted models (*P*<0.05-0.01; Table S5).

**Fig. 3.**
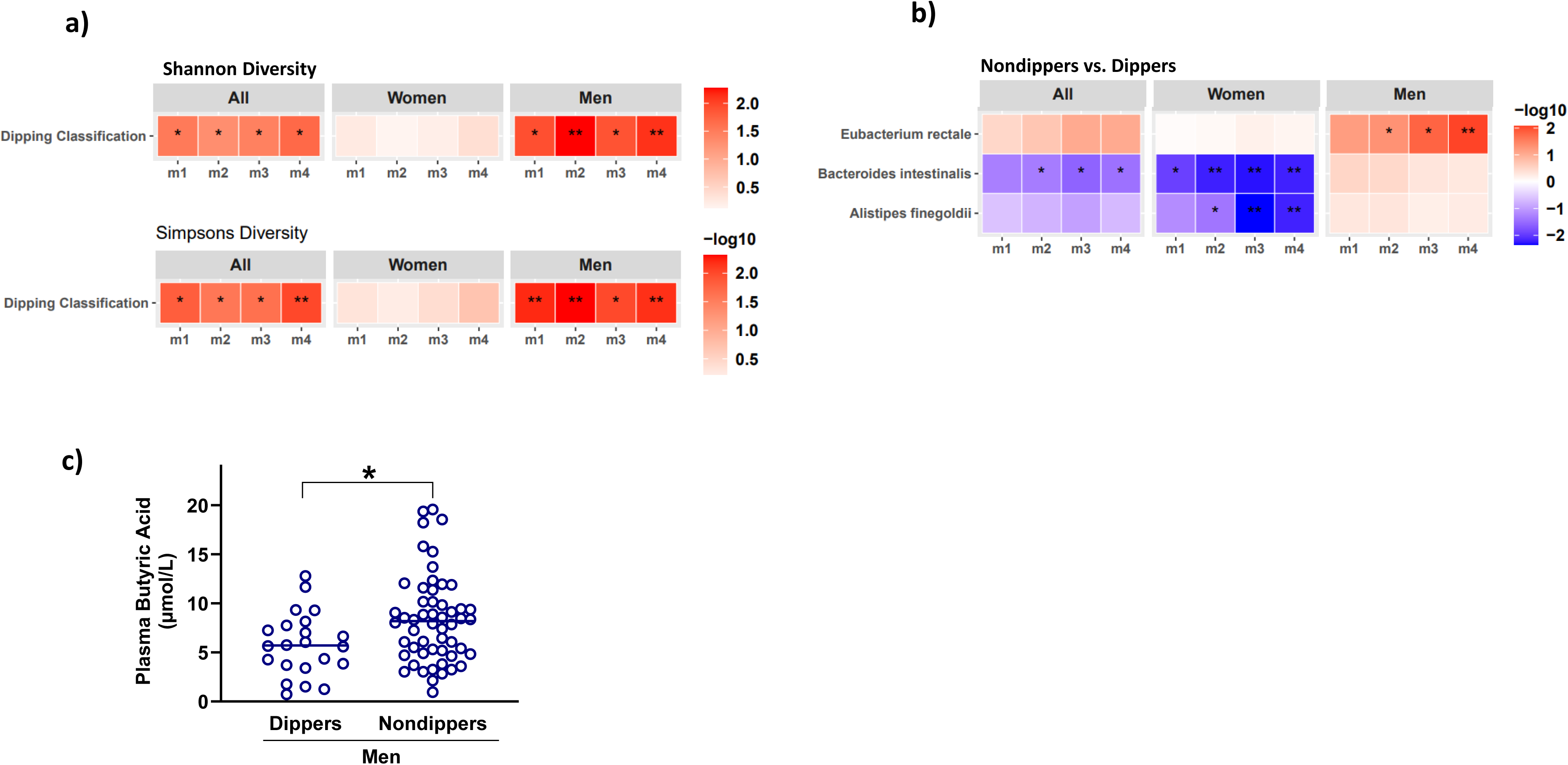
Gut microbiome and short-chain fatty acids associations with nighttime dipping in the whole cohort and sex-stratified analyses. (**a**) Association between Shannon diversity (top panel) and Simpson’s diversity (bottom panel) with nondippers vs. dippers. (**b**) Gut bacterial species associated with nondippers vs. dippers through linear regression analysis under 4 different models for covariate adjustment. Only species with at least < 0.01 significance in at least one model are shown. (**c**) Box-plots showing plasma butyric acid levels in dippers vs. nondipper men. Covatiate-adjusted models (m), m1: no covariate adjustment, m2: age; sex; BMI, m3: m2 + sodium intake based on spot urine analysis; serum glucose, triglyceride, HDL and LDL cholesterol; smoking and menopause status; liver steatosis by CAP score, m4: m3 +. 24-hour mean SBP or DBP and sleep latency. The statistically significant *p* values are indicated as \**P* < 0.05, \*\**P* < 0.01.

### Functional gut microbiome and BP variability

Sex-stratified analyses revealed significant associations between distinct functional genes and BP variability indices in men and women (Figs. S2-S3). The KEGG pathways “carbon fixation pathways in prokaryotes (ko00720) and “amino acid related enzymes (ko01007) were negatively associated with 24-hour SBP CoV in men (*P*<0.05-0.01 in m1-m4; Fig. S4, top panel). Higher 24-hour DBP CoV was associated with reduced “cytoskeleton proteins (ko04812)” and increased “transfer RNA biogenesis (ko03016)” in women (Fig. S2, bottom panel). In men with greater DBP CoV, “peptidoglycan biosynthesis and degradation proteins (ko01011)” pathway was significantly enriched (*P*<0.01-0.001 in m1-m4; Fig. S2, bottom panel). Furthermore, “signal transduction mechanisms” and “biotin metabolism (ko00780) pathways were significantly enriched in men with higher diastolic BP CoV (Fig. S2, bottom panel). Nondipping status was associated with a significant reduction of “bacterial toxin (ko02042)” genes in men (all *P*<0.05-0.01 in m1-m4; Fig. S3). Taken together, shotgun sequencing data revealed several previously unknown sex-linked associations between GM species, functional pathways, and GM metabolites with 24-hour BP variability indices in our cohort.

## Discussion

In this cross-sectional study, we demonstrate that GM and its metabolites are associated with 24-hour BP variability represented here as 24-hour SBP/DBP CoV and nighttime dipping in a sex-specific manner. To date, none but one clinical study has investigated the associations between 24-hour BP variability and GM profiled by 16S rRNA gene sequencing^12^, which lacks resolution for GM characterization at the species level as well as identifying low-abundance species^13^. In the present study, we identified GM species and SCFAs significantly associated with 24-hour BP variability indices despite adjusting for several confounding variables, such as blood glucose, LDL, and HDL, sodium intake, menopause status, sleep latency and mean 24-hour BP (Figure 4).

**Figure 4.**
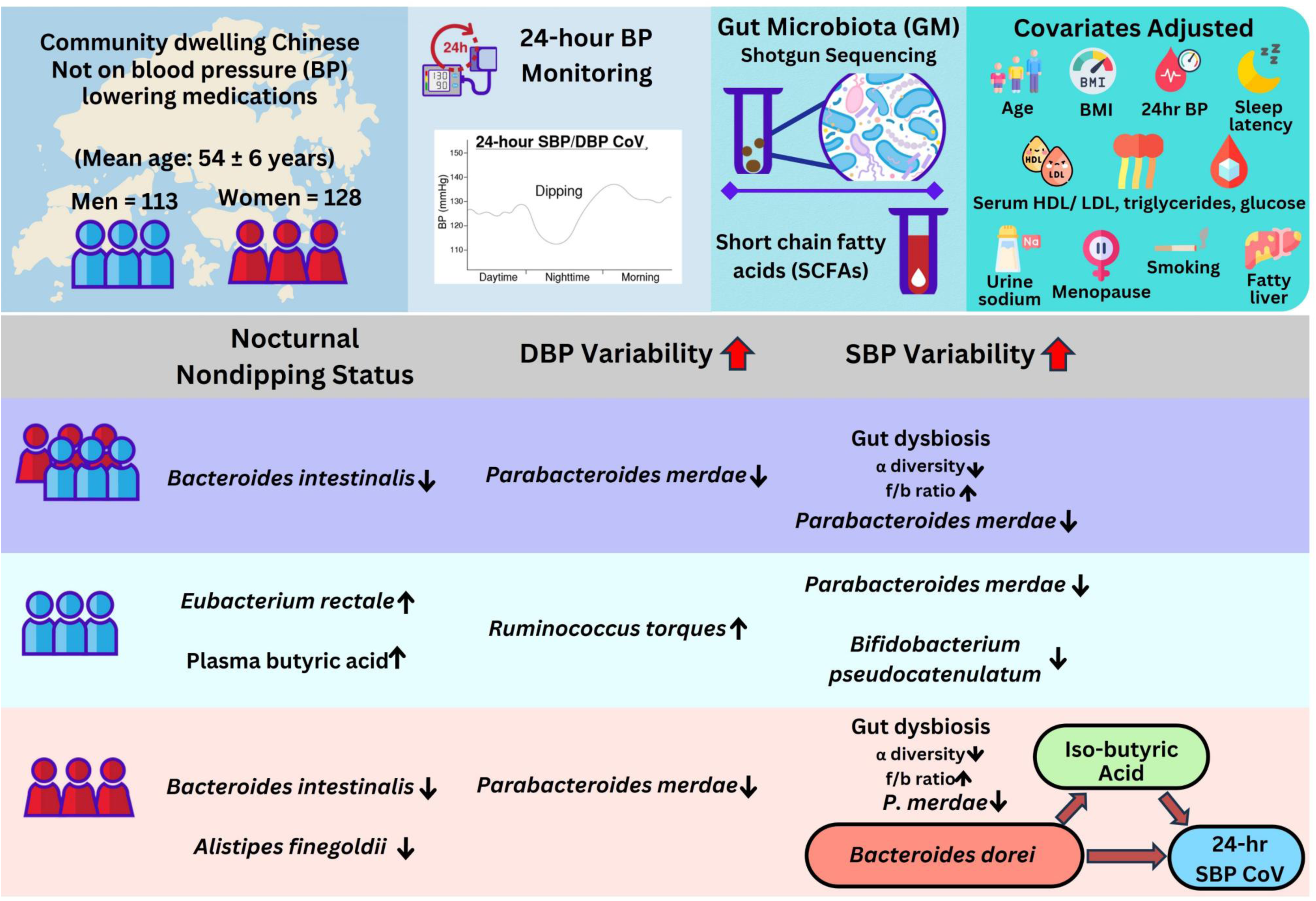
Graphical abstract: Sex-specific associations between gut microbiome and indices of 24-hour BP variability. BP, blood pressure; BMI, body mass index; CAP, controlled attenuation parameter; CoV, coefficient of variation; DBP, diastolic BP; GM, gut microbiota; HDL, high-density lipoprotein; LDL, low-density lipoprotein; SBP, systolic BP; SCFAs, short-chain fatty acids.

We recently reported that significant sex differences exist in GM associations with 24-hour BP and demonstrated that GM alterations were more significantly associated with hypertension in women than men^15^. Liang et al demonstrated that GM exhibited circadian rhythmicity over 24 hours, with more pronounced changes in female animal models compared to males, suggesting that sex may influence GM circadian oscillations^14^. Therefore, we asked if there are sex-linked differences in GM associations with 24-hour BP variability. GM α-diversity was significantly reduced in women with higher SBP CoV indicating reduced GM richness in women, but not men, with greater systolic BP variability in our cohort. Reduced α-diversity is a marker of GM dysbiosis and has been associated with CVD^7^. These results suggest that gut dysbiosis may contribute to the increased systolic BP variability, particularly in women. We further demonstrated that bacterial species known for health benefits, *Parabacteroides merdae*, *Bacteroides dorei* and *Bifidobactreium pseudocatenulatum*, had a significant inverse relationship with systolic and diastolic BP variability in both men and women. As shown in fig 2a and Table S4, the associations between these bacterial species and SBP variability remained significant after adjusting for various confounding variables and risk factors. *B. dorei* and *P. merdae* are noteworthy due to their known benefits on metabolic and cardiovascular health^30,31^. Oral administration of *B. dorei* and *P. merdae* have been reported to alleviate atherosclerotic lesions in animal models^30,31^. Taken together with our findings, these potentially beneficial bacterial taxa may provide novel gut microbiome interventions to alleviate systolic BP variability.

GM-derived SCFAs exhibit 24-hour oscillations, and influence host metabolic and inflammatory processes under circadian cues^32,33^. Thus, SCFAs may have a role in mediating 24-hour BP variability. An important component of BP variability, the nocturnal nondipping BP pattern, has been associated with cardiovascular events and organ damage^34^. In a small study, nondipper males (n=12) had a higher level of SCFAs, including butyric acid in the stool than dippers (n=26)^35^. In line with these findings, we detected significantly higher levels of plasma butyric acid and enrichment of a butyric acid-producer GM species, *Eubacterium rectale*^36^ in men with nondipping pattern in our cohort. Contrastingly, nondipping status in women was associated with a significantly reduced abundance of potentially beneficial commensal bacterial species, *Bacteroides intestinalis*^37^. These results indicate that GM may have distinct sex-linked mechanisms of mediating nocturnal dipping patterns. Furthermore, plasma iso-butyric acid had a negative association with 24-hour SBP CoV in the whole cohort. Most noteworthy, the negative correlation between iso-butyric acid and 24-hour SBP CoV was significant in women, but not in men, which persisted after adjusting for various confounding factors in sex-stratified analyses. Iso-butyric acid is a by-product of microbial fermentation of non-digestible proteins^38^. Although the biological functions and mechanisms of iso-butyric acid are not fully understood, a previous study reported that reduction in sodium intake decreased BP and increased serum concentrations of various SCFAs, including iso-butyric acid in an untreated mixed men-women cohort^39^, suggesting a potential role of iso-butyric acid in regulating BP. Furthermore, *B dorei*, which was found to have a significant negative association with 24-hour SBP CoV, had a significant positive correlation with plasma iso-butyric acid in women. Mediation analysis showed that *B dorei*’s association with 24-hour SBP CoV was mediated by plasma iso-butyric acid in women, which was significant after adjusting for various confounding variables and risk factors in statistical model 3, and was borderline significant after further adjusting for mean 24-hour SBP and sleep latency, as shown in Fig 2c/Table 2.

**Table 2.** Mediation of 24-hour SBP CoV by *Bacteroides dorei* via plasma iso-butyric acid in women. Legend. Casual mediation analysis was carried out using nonparametric bootstrap confidence intervals with the percentile method and 1000 simulations. Statistically significant *p* values are indicated as “*” 0.05, “†” P 0.01. Covariate-adjusted models, m3: age; sex; BMI; sodium intake based on spot urine analysis; serum glucose, triglyceride, HDL and LDL cholesterol; smoking and menopause status; hepatic steatosis CAP score; m4: m3 + 24-hour mean SBP and sleep latency. Abbreviations: ACME, average casual mediation effect; ADE, average direct effect; CAP, controlled attenuation parameter; CI, confidence intervals; HDL, high-density lipoprotein; LDL, low-density lipoprotein; SBP, systolic blood pressure.

| Mediation parameter | Model (m) | Estimate | 95% CI lower | 95% CI upper | P value |
| --- | --- | --- | --- | --- | --- |
| ACME | m3 | -1.6478 | -3.5990 | -0.19 | 0.028* |
|  | m4 | -1.41492 | -3.49478 | -0.03 | 0.048* |
| ADE | m3 | -4.0830 | -8.4848 | 0.20 | 0.064 |
|  | m4 | -4.63358 | -9.09325 | -0.08 | 0.050* |
| Total effect | m3 | -5.7308 | -9.8908 | -1.88 | 0.002† |
|  | m4 | -6.04849 | -10.07010 | -2.13 | 0.010† |
| Prop. mediated | m3 | 0.2875 | 0.0307 | 1.06 | 0.030* |
|  | m4 | 0.23393 | -0.00923 | 0.90 | 0.058 |

We acknowledge that the present study also has several limitations. First, our results are based on cross-sectional analysis and mechanistic studies in animal models need to be carried out to determine the causality. Second, our sample size, although larger than any previous study investigating the association between 24-hour BP variability and GM globally, is relatively small, and needs to be validated in larger multi-ethnicity cohorts. The present study also lacks analysis of other microbial kingdoms, including fungi, viruses and archaea, which can be identified using shotgun sequencing data. Despite these limitations, we have addressed several research gaps in the field by employing deep sequencing techniques to investigate the relationship between both SBP/DBP variability, while adjusting for various confounding factors, thus providing robust evidence for association between GM and 24-hour BP variability.

In summary, we demonstrated that higher systolic BP variability was associated with markers of gut dysbiosis and reduction in potentially beneficial gut bacteria, particularly in women, suggesting that the association between GM and 24-hour BP variability may be sex-linked (Figure 4). Furthermore, we identified several bacterial species which may be investigated for therapeutic potential to manage 24-hour BP variability, thus mitigating a significant risk factor for cardiovascular disease.

## Supplementary material

Supplementary figures S1-S3 and supplementary tables S1-S7

## Acknowledgments

The authors gratefully acknowledge Dr Ruby Szeto, Dr Isabelle Ngai, Dr Carmen Ho, Dr Cecilia Cheuk, Mr. Brian Shum, Mr. Sean Wong, Mr. Jacky Yueh, Mr. Jacky Wong, Mr. James Ng and Mr. Chapman for their assistance in data collection of this cohort. We thank the Genomics Core of the Centre for PanorOmic Sciences (CPOS), The University of Hong Kong for providing sample processing and metagenomic sequencing services.

## Author Contributions

KKL conceived and designed the study, and KKL, JWKH, PDV, GQ and KCT interpreted the data. PDV and GQ wrote the manuscript. MSSH, TKKTSP, CNMC, YHC, LKT, YHT, JWX, SSWL, RKCL, CPLL, CCWL, TLH and BYC were involved in the recruitment of participants, clinical assessments, food questionnaire logging, and stool, urine and blood sample collection. GQ conducted the bioinformatics analysis supervised by JWKH. PDV, CPLL, SSWL conducted experiments to estimate serum biomarkers and urine sodium. KSL and EKKL carried out the stool and plasma SCFA estimation supervised by HE-N. CPLL, LKT and TKKTSP analyzed dietary food intake supervised by JCYL. S.Y.L. provided the infrastructure for and supervised the metagenomic sequencing. KKL, JWKH, HE-N, SYL, IFNH, MFY contributed to financial resources. All authors analyzed the data, discussed the results, commented on the manuscript, and approved the final version of the manuscript.

## Funding

This work was supported in part by the Croucher Foundation, the Hong Kong Jockey Club Charities Trust, and AIR@InnoHK administered by Innovation and Technology Commission.

## Conflict of interest

KKL received grants from Research Fund Secretariat of the Food and Health Bureau, Innovation and Technology Bureau, Research Grants Council, Amgen, Boehringer Ingelheim, Eisai and Pfizer; and consultation fees from Amgen, Boehringer Ingelheim, Daiichi Sankyo and Sanofi, all outside the submitted work. SYL has received research sponsorship from Pfizer, Merck, Servier and Curegenix, all outside the submitted work.

## Data Availability

All of the processed GM species abundance data and clinical metadata, the scripts for data processing and generating figures have been deposited in an HKU GitHub repository which can be accessed using the following link: https://github.com/holab-hku/bp_variability_study. Raw data are available from the corresponding author upon reasonable request.

## Notes

### Competing Interest Statement

The authors have declared no competing interest.

