## Supplementary material for "Associations Between Gut Microbiome and 24-Hour Blood Pressure Variability: A Cross-sectional Study Highlighting Sex Differences and Potential Therapeutic Targets"

[1]Department of Medicine, School of Clinical Medicine, Li Ka Shing (LKS) Faculty of Medicine, The University of Hong Kong, Hong Kong Special Administrative Region (SAR), China; [2]State Key Laboratory of Brain and Cognitive Sciences, The University of Hong Kong, Hong Kong, China; [3]School of Biomedical Sciences, LKS Faculty of Medicine, The University of Hong Kong, Hong Kong SAR, China; [4]Laboratory of Data Discovery for Health Limited (D$^2$4H), Hong Kong Science Park, Hong Kong S.A.R., China; [5]School of Biological Sciences, Faculty of Science, The University of Hong Kong, Hong Kong SAR, China; [6]Department of Pathology, School of Clinical Medicine, The University of Hong Kong, Queen Mary Hospital, Pokfulam, Hong Kong SAR, China; [7]The Jockey Club Centre for Clinical Innovation and Discovery, LKS Faculty of Medicine, The University of Hong Kong, Pokfulam, Hong Kong SAR,

China; [8]Centre for PanorOmic Sciences, LKS Faculty of Medicine, The University of Hong Kong, Pokfulam, Hong Kong SAR, China; [9]Institute of Public Health and Clinical Nutrition, University of Eastern Finland, Kuopio 70211, Finland; [10]Discipline of Dietetics, Department of Nursing and Allied Health, School of Health Sciences, Swinburne University of Technology, Melbourne, Australia.

#PDV and GQ contributed equally to this work.

#### Supplementary Figures

**Fig. S1**

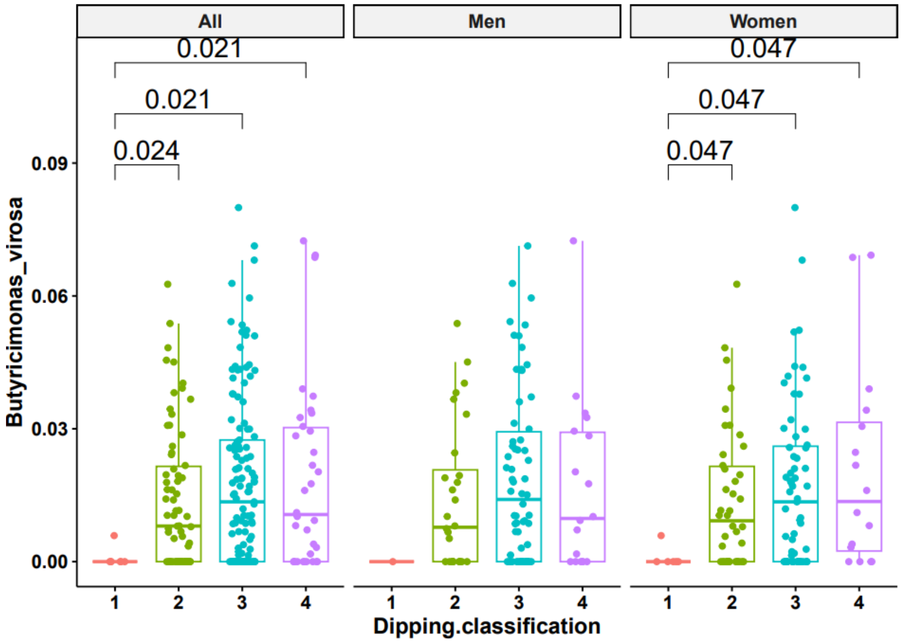


**Fig. S1. Association of GM species with dipping status.** Box plots showing associations of Butyricimonas Virosa with dipping status in the whole cohort, men and women. Dipping classification: 1, extreme dippers; 2, dippers; 3, nondippers; 4, reverse dippers. The statistical significance was determined by unpaired t-test: $^{**}P < 0.01$, $^{***}P < 0.001$.

Fig S2

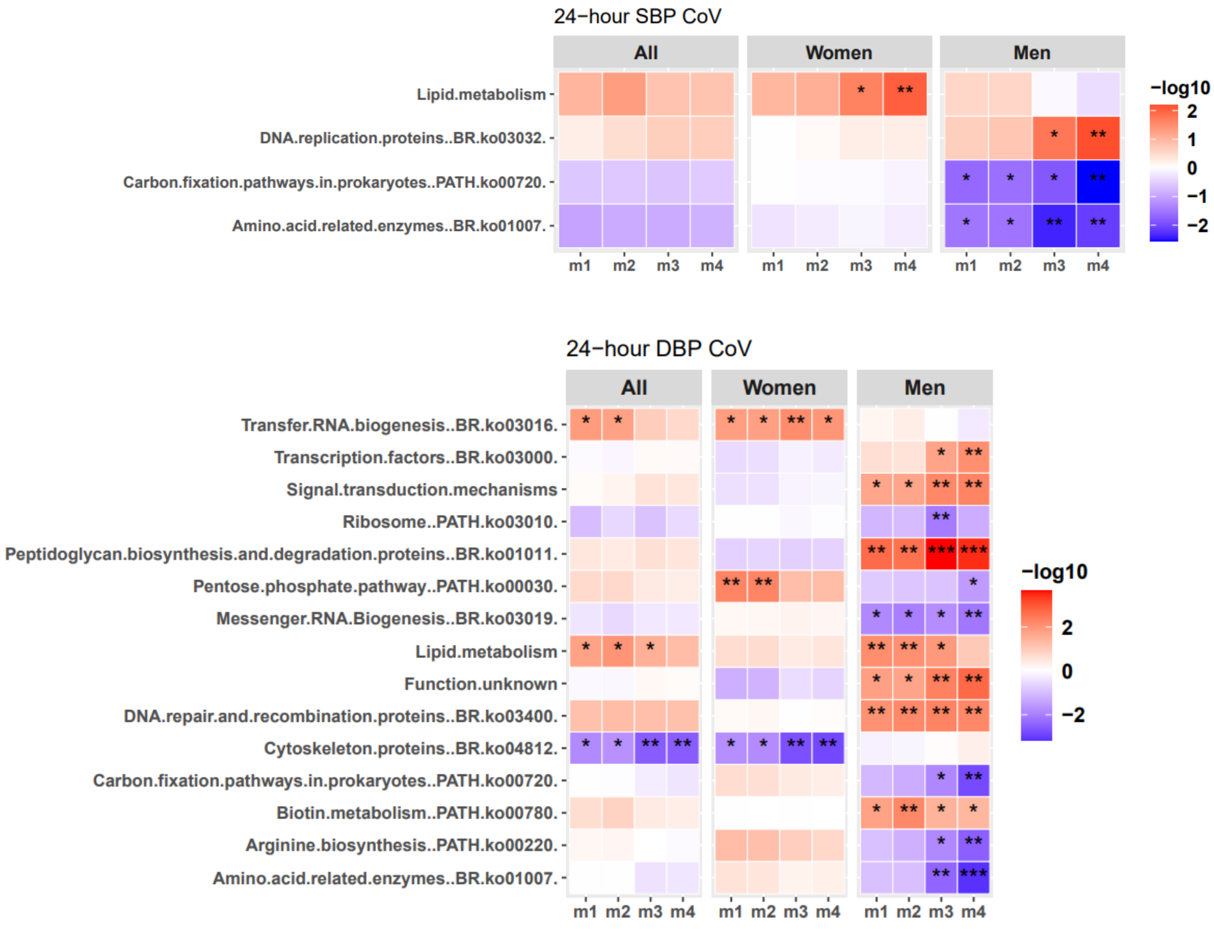


**Fig. S2. Association of KEGG functional pathways with 24-hour BP variability.** Association of KEGG functional pathways with 24-hour SBP (top panel) / DBP (bottom panel) CoV. All analyses were performed under four different linear regression models for covariate adjustment, m1: no covariate adjustment, m2: age; sex; BMI, m3: m2 + sodium intake based on spot urine analysis; serum glucose, triglyceride, HDL and LDL cholesterol; smoking and menopause status; liver steatosis by CAP score, m4: m3 +. 24-hour mean SBP or DBP and sleep latency. The statistically significant p values are indicated as *P < 0.05, **P < 0.01, ***P < 0.001

**Fig S3**

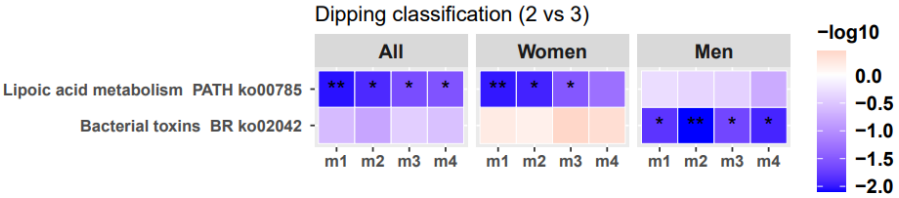


**Fig. S3. Association of KEGG functional pathways with BP variability indices.** Association of KEGG functional pathways with nondipping status (3) vs dippers (2). Covariate adjusted models, m1: no covariate adjustment, m2: age; sex; BMI, m3: m2 + sodium intake based on spot urine analysis; serum glucose, triglyceride, HDL and LDL cholesterol; smoking and menopause status; liver steatosis by CAP score, m4: m3 +. 24-hour mean SBP or DBP and sleep latency. The statistically significant p values are indicated as *P < 0.05, **P < 0.01, ***P < 0.001.

### Supplementary Tables

**Table S1. Sex differences in BP variability**

| Diurnal BP variability index | Model (m) | *P*-value | Coefficient | Standard error |
|---|---|---|---|---|
| 24-hour SBP CoV | m1 | 0.001 | 1.115 | 0.318 |
| | m2 | 0.000 | 1.279 | 0.339 |
| | m3 | 0.002 | 1.340 | 0.433 |
| | m4 | 0.001 | 1.475 | 0.446 |
| Daytime SBP CoV | m1 | 0.000 | 1.124 | 0.313 |
| | m2 | 0.001 | 1.110 | 0.333 |
| | m3 | 0.001 | 1.380 | 0.423 |
| | m4 | 0.001 | 1.442 | 0.438 |
| Nighttime SBP CoV | m1 | 0.484 | 0.327 | 0.467 |
| | m2 | 0.090 | 0.838 | 0.492 |
| | m3 | 0.493 | 0.430 | 0.626 |
| | m4 | 0.313 | 0.650 | 0.642 |
| 24-hour DBP CoV | m1 | 0.002 | 1.444 | 0.465 |
| | m2 | 0.018 | 1.190 | 0.498 |
| | m3 | 0.009 | 1.696 | 0.643 |
| | m4 | 0.038 | 1.398 | 0.670 |
| Daytime DBP CoV | m1 | 0.000 | 1.879 | 0.495 |
| | m2 | 0.006 | 1.457 | 0.527 |
| | m3 | 0.002 | 2.192 | 0.683 |
| | m4 | 0.012 | 1.804 | 0.707 |
| Nighttime DBP CoV | m1 | 0.366 | -0.526 | 0.582 |
| | m2 | 0.863 | -0.107 | 0.620 |
| | m3 | 0.435 | -0.611 | 0.781 |
| | m4 | 0.179 | -1.112 | 0.824 |

Legend. Logistic regression model results of differences in BP variability indices among men and women. Four models for covariate adjustment were used to control for confounding factors, m1: no covariate adjustment, m2: age; sex; BMI, m3: m2 + sodium intake based on spot urine analysis; serum glucose, triglyceride, HDL, and LDL cholesterol; smoking status; menopause status; liver steatosis by CAP score, m4: m3 + 24-hour mean SBP or DBP; sleep latency. Abbreviations: BP, blood pressure; CoV, coefficient of variation; DBP, diastolic blood pressure; HDL, high-density lipoprotein; LDL, low-density lipoprotein; SBP, systolic blood pressure.

**Table S2. Shannon index associations with 24-hour BP variability indices.**

| BP variability index | Sex | Model (m) | *P*-value | Coefficient | Standard error |
|---|---|---|---|---|---|
| 24-hour SBP CoV | All | m1 | 0.114 | -0.013 | 0.008 |
| | | m2 | 0.150 | -0.013 | 0.009 |
| | | m3 | 0.027 | -0.022 | 0.010 |
| | | m4 | 0.022 | -0.023 | 0.010 |
| | Men | m1 | 0.850 | 0.003 | 0.017 |
| | | m2 | 0.900 | 0.002 | 0.017 |
| | | m3 | 0.824 | 0.005 | 0.022 |
| | | m4 | 0.926 | 0.002 | 0.023 |
| | Women | m1 | 0.075 | -0.018 | 0.010 |
| | | m2 | 0.108 | -0.017 | 0.010 |
| | | m3 | 0.016 | -0.028 | 0.012 |
| | | m4 | 0.022 | -0.028 | 0.012 |
| Daytime SBP CoV | All | m1 | 0.452 | -0.006 | 0.009 |
| | | m2 | 0.499 | -0.006 | 0.009 |
| | | m3 | 0.159 | -0.014 | 0.010 |
| | | m4 | 0.175 | -0.014 | 0.010 |
| | Men | m1 | 0.270 | 0.016 | 0.015 |
| | | m2 | 0.287 | 0.016 | 0.015 |
| | | m3 | 0.253 | 0.022 | 0.019 |
| | | m4 | 0.197 | 0.026 | 0.020 |
| | Women | m1 | 0.141 | -0.016 | 0.011 |
| | | m2 | 0.125 | -0.017 | 0.011 |
| | | m3 | 0.024 | -0.028 | 0.012 |
| | | m4 | 0.031 | -0.028 | 0.013 |
| Nighttime SBP CoV | All | m1 | 0.332 | -0.006 | 0.006 |
| | | m2 | 0.427 | -0.005 | 0.006 |
| | | m3 | 0.406 | -0.006 | 0.007 |
| | | m4 | 0.312 | -0.007 | 0.007 |
| | Men | m1 | 0.725 | 0.003 | 0.009 |
| | | m2 | 0.743 | 0.003 | 0.009 |
| | | m3 | 0.353 | 0.011 | 0.011 |
| | | m4 | 0.418 | 0.010 | 0.012 |
| | Women | m1 | 0.151 | -0.011 | 0.008 |
| | | m2 | 0.349 | -0.008 | 0.008 |
| | | m3 | 0.197 | -0.012 | 0.009 |
| | | m4 | 0.240 | -0.011 | 0.010 |
| 24-hour DBP CoV | All | m1 | 0.150 | -0.008 | 0.006 |
| | | m2 | 0.147 | -0.009 | 0.006 |
| | | m3 | 0.073 | -0.012 | 0.007 |

| | | | | | |
|---|---|---|---|---|---|
| | | m4 | 0.049 | -0.014 | 0.007 |
| | Men | m1 | 0.757 | 0.003 | 0.009 |
| | | m2 | 0.642 | 0.004 | 0.010 |
| | | m3 | 0.434 | 0.009 | 0.011 |
| | | m4 | 0.663 | 0.005 | 0.012 |
| | Women | m1 | 0.060 | -0.014 | 0.008 |
| | | m2 | 0.048 | -0.015 | 0.007 |
| | | m3 | 0.061 | -0.017 | 0.009 |
| | | m4 | 0.075 | -0.017 | 0.009 |
| Daytime DBP CoV | All | m1 | 0.281 | -0.006 | 0.005 |
| | | m2 | 0.260 | -0.006 | 0.006 |
| | | m3 | 0.199 | -0.008 | 0.006 |
| | | m4 | 0.159 | -0.009 | 0.006 |
| | Men | m1 | 0.455 | 0.007 | 0.009 |
| | | m2 | 0.376 | 0.008 | 0.009 |
| | | m3 | 0.224 | 0.012 | 0.010 |
| | | m4 | 0.323 | 0.011 | 0.011 |
| | Women | m1 | 0.080 | -0.013 | 0.007 |
| | | m2 | 0.042 | -0.015 | 0.007 |
| | | m3 | 0.094 | -0.015 | 0.009 |
| | | m4 | 0.109 | -0.015 | 0.009 |
| Nighttime DBP CoV | All | m1 | 0.628 | -0.002 | 0.005 |
| | | m2 | 0.688 | -0.002 | 0.005 |
| | | m3 | 0.500 | -0.004 | 0.005 |
| | | m4 | 0.457 | -0.004 | 0.006 |
| | Men | m1 | 0.424 | 0.005 | 0.006 |
| | | m2 | 0.390 | 0.006 | 0.006 |
| | | m3 | 0.239 | 0.009 | 0.008 |
| | | m4 | 0.284 | 0.009 | 0.008 |
| | Women | m1 | 0.143 | -0.010 | 0.007 |
| | | m2 | 0.319 | -0.007 | 0.007 |
| | | m3 | 0.288 | -0.009 | 0.008 |
| | | m4 | 0.295 | -0.009 | 0.009 |

Legend. Linear regression analyses of association between Shannon diversity and 24-hour BP variability indices. Four models for covariate adjustment were used to control for confounding factors, m1: no covariate adjustment, m2: age; sex; BMI, m3: m2 + sodium intake based on spot urine analysis; serum glucose, triglyceride, HDL, and LDL cholesterol; smoking status; menopause status; liver steatosis by CAP score, m4: m3 + 24-hour mean SBP or DBP; sleep latency. Abbreviations: BP, blood pressure; CoV, coefficient of variation; DBP, diastolic blood pressure; HDL, high-density lipoprotein; LDL, low-density lipoprotein; SBP, systolic blood pressure.

**Table S3. Associations between Firmicutes/Bacteroidetes (f/b) ratio and 24-hour BP variability indices.**

| BP variability index | Sex | Model (m) | *P*-value | Coefficient | Standard error |
|---|---|---|---|---|---|
| 24-hour SBP CoV | All | m1 | 0.060 | 0.224 | 0.118 |
| | | m2 | 0.092 | 0.206 | 0.122 |
| | | m3 | 0.015 | 0.318 | 0.129 |
| | | m4 | 0.044 | 0.252 | 0.124 |
| | Men | m1 | 0.286 | 0.206 | 0.192 |
| | | m2 | 0.188 | 0.249 | 0.188 |
| | | m3 | 0.076 | 0.484 | 0.269 |
| | | m4 | 0.372 | 0.205 | 0.229 |
| | Women | m1 | 0.242 | 0.190 | 0.162 |
| | | m2 | 0.281 | 0.173 | 0.160 |
| | | m3 | 0.049 | 0.303 | 0.152 |
| | | m4 | 0.063 | 0.301 | 0.160 |
| Daytime SBP CoV | All | m1 | 0.850 | 0.023 | 0.121 |
| | | m2 | 0.923 | 0.012 | 0.125 |
| | | m3 | 0.524 | 0.086 | 0.134 |
| | | m4 | 0.599 | 0.067 | 0.128 |
| | Men | m1 | 0.584 | -0.095 | 0.172 |
| | | m2 | 0.709 | -0.063 | 0.169 |
| | | m3 | 0.861 | -0.042 | 0.238 |
| | | m4 | 0.665 | -0.085 | 0.196 |
| | Women | m1 | 0.921 | 0.017 | 0.174 |
| | | m2 | 0.723 | 0.061 | 0.172 |
| | | m3 | 0.292 | 0.173 | 0.163 |
| | | m4 | 0.325 | 0.169 | 0.170 |
| Nighttime SBP CoV | All | m1 | 0.017 | 0.198 | 0.082 |
| | | m2 | 0.022 | 0.192 | 0.083 |
| | | m3 | 0.033 | 0.193 | 0.090 |
| | | m4 | 0.073 | 0.156 | 0.086 |
| | Men | m1 | 0.440 | 0.081 | 0.104 |
| | | m2 | 0.420 | 0.083 | 0.102 |
| | | m3 | 0.461 | 0.105 | 0.143 |
| | | m4 | 0.796 | 0.030 | 0.116 |
| | Women | m1 | 0.031 | 0.264 | 0.121 |
| | | m2 | 0.095 | 0.211 | 0.126 |
| | | m3 | 0.132 | 0.181 | 0.119 |
| | | m4 | 0.164 | 0.177 | 0.126 |
| 24-hour DBP CoV | All | m1 | 0.309 | 0.083 | 0.082 |
| | | m2 | 0.419 | 0.067 | 0.083 |

| | | | | | |
|---|---|---|---|---|---|
| | | m3 | 0.030 | 0.191 | 0.087 |
| | | m4 | 0.089 | 0.147 | 0.086 |
| | Men | m1 | 0.081 | 0.188 | 0.107 |
| | | m2 | 0.201 | 0.138 | 0.107 |
| | | m3 | 0.085 | 0.235 | 0.135 |
| | | m4 | 0.172 | 0.164 | 0.119 |
| | Women | m1 | 0.931 | -0.011 | 0.121 |
| | | m2 | 0.952 | 0.007 | 0.118 |
| | | m3 | 0.159 | 0.169 | 0.119 |
| | | m4 | 0.211 | 0.157 | 0.125 |
| Daytime DBP CoV | All | m1 | 0.812 | -0.018 | 0.076 |
| | | m2 | 0.667 | -0.034 | 0.079 |
| | | m3 | 0.475 | 0.060 | 0.083 |
| | | m4 | 0.654 | 0.037 | 0.082 |
| | Men | m1 | 0.712 | 0.037 | 0.100 |
| | | m2 | 0.973 | -0.003 | 0.099 |
| | | m3 | 0.565 | 0.071 | 0.123 |
| | | m4 | 0.708 | 0.040 | 0.106 |
| | Women | m1 | 0.392 | -0.098 | 0.114 |
| | | m2 | 0.645 | -0.052 | 0.113 |
| | | m3 | 0.634 | 0.054 | 0.114 |
| | | m4 | 0.745 | 0.039 | 0.121 |
| Nighttime DBP CoV | All | m1 | 0.005 | 0.185 | 0.066 |
| | | m2 | 0.007 | 0.179 | 0.066 |
| | | m3 | 0.015 | 0.176 | 0.072 |
| | | m4 | 0.052 | 0.136 | 0.070 |
| | Men | m1 | 0.020 | 0.170 | 0.072 |
| | | m2 | 0.034 | 0.152 | 0.071 |
| | | m3 | 0.177 | 0.134 | 0.098 |
| | | m4 | 0.187 | 0.107 | 0.081 |
| | Women | m1 | 0.051 | 0.213 | 0.108 |
| | | m2 | 0.190 | 0.145 | 0.110 |
| | | m3 | 0.281 | 0.119 | 0.109 |
| | | m4 | 0.312 | 0.116 | 0.114 |
| | | m4 | 0.619 | -0.023 | 0.047 |

Legend. Linear regression analyses of association between f/b ratio and 24-hour BP variability indices. Four models for covariate adjustment were used to control for confounding factors, m1: no covariate adjustment, m2: age; sex; BMI, m3: m2 + sodium intake based on spot urine analysis; serum glucose, triglyceride, HDL, and LDL cholesterol; smoking status; menopause status; liver steatosis by CAP score, m4: m3 + 24-hour mean SBP or DBP; sleep latency. Abbreviations: BP, blood pressure; CoV, coefficient of variation; DBP, diastolic blood pressure; HDL, high-density lipoprotein; LDL, low-density lipoprotein; SBP, systolic blood pressure.

**Table S4. GM species associated with 24-hour BP variability indices**

| BP variability index | GM species | Sex | Model (m) | *P*-value | Coefficient | Standard error |
|---|---|---|---|---|---|---|
| 24-hour SBP CoV | *Parabacteroides merdae* | All | m1 | 0.008 | -11.776 | 4.419 |
| | | | m2 | 0.009 | -11.386 | 4.342 |
| | | | m3 | 0.001 | -14.767 | 4.558 |
| | | | m4 | 0.001 | -15.110 | 4.656 |
| | | Men | m1 | 0.047 | -9.709 | 4.833 |
| | | | m2 | 0.050 | -9.660 | 4.863 |
| | | | m3 | 0.006 | -12.539 | 4.445 |
| | | | m4 | 0.006 | -12.275 | 4.316 |
| | | Women | m1 | 0.045 | -13.689 | 6.746 |
| | | | m2 | 0.076 | -12.382 | 6.914 |
| | | | m3 | 0.028 | -17.031 | 7.624 |
| | | | m4 | 0.026 | -18.006 | 7.930 |
| | *Bacteroides dorei* | All | m1 | 0.176 | -2.293 | 1.689 |
| | | | m2 | 0.036 | -3.488 | 1.658 |
| | | | m3 | 0.193 | -2.295 | 1.758 |
| | | | m4 | 0.232 | -2.159 | 1.801 |
| | | Men | m1 | 0.445 | 1.502 | 1.958 |
| | | | m2 | 0.467 | 1.439 | 1.973 |
| | | | m3 | 0.161 | 2.550 | 1.804 |
| | | | m4 | 0.109 | 2.844 | 1.751 |
| | | Women | m1 | 0.014 | -6.177 | 2.487 |
| | | | m2 | 0.008 | -6.690 | 2.473 |
| | | | m3 | 0.029 | -6.082 | 2.743 |
| | | | m4 | 0.015 | -7.127 | 2.871 |
| | *Bifidobacterium pseudocatenulatum* | All | m1 | 0.773 | 0.322 | 1.112 |
| | | | m2 | 0.982 | -0.024 | 1.086 |
| | | | m3 | 0.675 | 0.479 | 1.140 |
| | | | m4 | 0.755 | 0.363 | 1.163 |
| | | Men | m1 | 0.005 | -3.703 | 1.279 |
| | | | m2 | 0.006 | -3.629 | 1.302 |
| | | | m3 | 0.076 | -2.227 | 1.240 |
| | | | m4 | 0.024 | -2.769 | 1.205 |
| | | Women | m1 | 0.222 | 1.984 | 1.615 |
| | | | m2 | 0.256 | 1.840 | 1.611 |
| | | | m3 | 0.375 | 1.664 | 1.868 |
| | | | m4 | 0.344 | 1.835 | 1.929 |
| 24-hour DBP CoV | *Parabacteroides merdae* | All | m1 | 0.033 | -13.884 | 6.466 |
| | | | m2 | 0.014 | -15.815 | 6.393 |
| | | | m3 | 0.004 | -19.977 | 6.802 |
| | | | m4 | 0.003 | -20.697 | 6.782 |

| | | | | | | |
|---|---|---|---|---|---|---|
| | | Men | m1 | 0.163 | -12.236 | 8.709 |
| | | | m2 | 0.151 | -12.448 | 8.617 |
| | | | m3 | 0.168 | -12.834 | 9.225 |
| | | | m4 | 0.164 | -12.087 | 8.595 |
| | | Women | m1 | 0.088 | -15.623 | 9.098 |
| | | | m2 | 0.063 | -17.637 | 9.403 |
| | | | m3 | 0.020 | -23.212 | 9.829 |
| | | | m4 | 0.023 | -23.735 | 10.265 |
| | *Ruminococcus torques* | All | m1 | 0.418 | 1.766 | 2.179 |
| | | | m2 | 0.271 | 2.373 | 2.150 |
| | | | m3 | 0.258 | 2.765 | 2.437 |
| | | | m4 | 0.242 | 2.930 | 2.495 |
| | | Men | m1 | 0.007 | 7.121 | 2.609 |
| | | | m2 | 0.006 | 7.207 | 2.595 |
| | | | m3 | 0.011 | 7.726 | 2.985 |
| | | | m4 | 0.011 | 7.679 | 2.930 |
| | | Women | m1 | 0.419 | -2.683 | 3.308 |
| | | | m2 | 0.446 | -2.558 | 3.342 |
| | | | m3 | 0.712 | -1.370 | 3.706 |
| | | | m4 | 0.920 | -0.406 | 4.007 |
| | *Clostridium sp CAG 58* | All | m1 | 0.049 | -17.231 | 8.723 |
| | | | m2 | 0.062 | -16.127 | 8.608 |
| | | | m3 | 0.010 | -26.022 | 9.951 |
| | | | m4 | 0.007 | -27.403 | 10.118 |
| | | Men | m1 | 0.630 | 5.310 | 10.984 |
| | | | m2 | 0.573 | 6.159 | 10.886 |
| | | | m3 | 0.770 | -4.188 | 14.291 |
| | | | m4 | 0.334 | -13.013 | 13.383 |
| | | Women | m1 | 0.005 | -36.469 | 12.774 |
| | | | m2 | 0.004 | -37.455 | 12.920 |
| | | | m3 | 0.018 | -34.499 | 14.310 |
| | | | m4 | 0.021 | -35.593 | 15.175 |
| | *Clostridium bolteae* | All | m1 | 0.149 | -21.199 | 14.642 |
| | | | m2 | 0.131 | -21.789 | 14.384 |
| | | | m3 | 0.269 | -17.169 | 15.491 |
| | | | m4 | 0.366 | -13.991 | 15.449 |
| | | Men | m1 | 0.367 | 16.818 | 18.571 |
| | | | m2 | 0.307 | 18.906 | 18.433 |
| | | | m3 | 0.247 | 23.945 | 20.534 |
| | | | m4 | 0.454 | 14.813 | 19.678 |
| | | Women | m1 | 0.008 | -56.662 | 21.158 |
| | | | m2 | 0.007 | -58.881 | 21.422 |
| | | | m3 | 0.183 | -32.119 | 23.930 |

| | | | | | | |
|---|---|---|---|---|---|---|
| | | | m4 | 0.278 | -28.357 | 25.971 |
| | *Acidaminococcus intestini* | All | m1 | 0.038 | 10.818 | 5.174 |
| | | | m2 | 0.023 | 11.856 | 5.180 |
| | | | m3 | 0.201 | 7.608 | 5.931 |
| | | | m4 | 0.387 | 5.133 | 5.916 |
| | | Men | m1 | 0.001 | 20.425 | 6.223 |
| | | | m2 | 0.004 | 19.034 | 6.412 |
| | | | m3 | 0.098 | 14.616 | 8.720 |
| | | | m4 | 0.181 | 11.220 | 8.304 |
| | | Women | m1 | 0.645 | 3.624 | 7.848 |
| | | | m2 | 0.581 | 4.441 | 8.026 |
| | | | m3 | 0.439 | 6.584 | 8.480 |
| | | | m4 | 0.562 | 5.041 | 8.658 |
| Daytime SBP CoV | *Bacteroides dorei* | All | m1 | 0.219 | -2.050 | 1.664 |
| | | | m2 | 0.043 | -3.320 | 1.630 |
| | | | m3 | 0.156 | -2.442 | 1.716 |
| | | | m4 | 0.144 | -2.591 | 1.767 |
| | | Men | m1 | 0.608 | 1.131 | 2.198 |
| | | | m2 | 0.646 | 1.017 | 2.212 |
| | | | m3 | 0.269 | 2.324 | 2.090 |
| | | | m4 | 0.265 | 2.324 | 2.072 |
| | | Women | m1 | 0.018 | -5.558 | 2.328 |
| | | | m2 | 0.009 | -6.169 | 2.311 |
| | | | m3 | 0.037 | -5.508 | 2.599 |
| | | | m4 | 0.017 | -6.642 | 2.735 |
| | *Bacteroides stercoris* | All | m1 | 0.013 | -4.659 | 1.869 |
| | | | m2 | 0.073 | -3.353 | 1.862 |
| | | | m3 | 0.011 | -5.131 | 1.986 |
| | | | m4 | 0.010 | -5.324 | 2.036 |
| | | Men | m1 | 0.044 | -4.280 | 2.099 |
| | | | m2 | 0.056 | -4.103 | 2.127 |
| | | | m3 | 0.025 | -4.816 | 2.106 |
| | | | m4 | 0.038 | -4.463 | 2.115 |
| | | Women | m1 | 0.404 | -2.603 | 3.112 |
| | | | m2 | 0.340 | -2.965 | 3.094 |
| | | | m3 | 0.071 | -6.376 | 3.490 |
| | | | m4 | 0.036 | -7.651 | 3.596 |
| | *Paraprevotella xylaniphila* | All | m1 | 0.095 | -9.220 | 5.497 |
| | | | m2 | 0.108 | -8.643 | 5.355 |
| | | | m3 | 0.010 | -14.753 | 5.639 |
| | | | m4 | 0.011 | -14.876 | 5.814 |
| | | Men | m1 | 0.423 | 5.247 | 6.518 |
| | | | m2 | 0.463 | 4.861 | 6.598 |

| | | | | | | |
|---|---|---|---|---|---|---|
| | | | m3 | 0.702 | -2.654 | 6.900 |
| | | | m4 | 0.889 | -0.961 | 6.878 |
| | | Women | m1 | 0.011 | -21.028 | 8.168 |
| | | | m2 | 0.016 | -19.875 | 8.153 |
| | | | m3 | 0.010 | -22.599 | 8.657 |
| | | | m4 | 0.021 | -21.652 | 9.232 |
| | *Alistipes indistinctus* | All | m1 | 0.301 | 8.328 | 8.036 |
| | | | m2 | 0.585 | 4.371 | 8.001 |
| | | | m3 | 0.543 | 5.107 | 8.379 |
| | | | m4 | 0.488 | 5.960 | 8.586 |
| | | Men | m1 | 0.042 | 21.750 | 10.547 |
| | | | m2 | 0.058 | 20.829 | 10.850 |
| | | | m3 | 0.008 | 29.145 | 10.728 |
| | | | m4 | 0.006 | 29.834 | 10.590 |
| | | Women | m1 | 0.706 | -4.265 | 11.292 |
| | | | m2 | 0.700 | -4.470 | 11.580 |
| | | | m3 | 0.609 | -6.437 | 12.550 |
| | | | m4 | 0.843 | -2.610 | 13.171 |
| Daytime DBP CoV | *Acidaminococcus intestini* | All | m1 | 0.043 | 11.347 | 5.564 |
| | | | m2 | 0.015 | 13.386 | 5.476 |
| | | | m3 | 0.201 | 8.072 | 6.291 |
| | | | m4 | 0.444 | 4.790 | 6.247 |
| | | Men | m1 | 0.000 | 24.078 | 6.652 |
| | | | m2 | 0.001 | 24.016 | 6.858 |
| | | | m3 | 0.034 | 20.649 | 9.568 |
| | | | m4 | 0.077 | 16.799 | 9.356 |
| | | Women | m1 | 0.833 | 1.750 | 8.296 |
| | | | m2 | 0.670 | 3.595 | 8.409 |
| | | | m3 | 0.661 | 3.940 | 8.964 |
| | | | m4 | 0.850 | 1.706 | 8.987 |
| | *Megasphaera elsdenii* | All | m1 | 0.306 | 9.953 | 9.692 |
| | | | m2 | 0.068 | 17.435 | 9.527 |
| | | | m3 | 0.189 | 14.158 | 10.743 |
| | | | m4 | 0.079 | 20.655 | 11.691 |
| | | Men | m1 | 0.051 | 23.224 | 11.790 |
| | | | m2 | 0.029 | 26.758 | 12.086 |
| | | | m3 | 0.043 | 34.575 | 16.776 |
| | | | m4 | 0.006 | 45.559 | 15.980 |
| | | Women | m1 | 0.681 | 6.086 | 14.754 |
| | | | m2 | 0.608 | 7.593 | 14.751 |
| | | | m3 | 0.862 | -2.627 | 15.113 |
| | | | m4 | 0.743 | 5.967 | 18.177 |
| | *Roseburia intestinalis* | All | m1 | 0.013 | -9.514 | 3.801 |

| | | | | | | |
|---|---|---|---|---|---|---|
| | | | m2 | 0.028 | -8.182 | 3.710 |
| | | | m3 | 0.095 | -7.043 | 4.200 |
| | | | m4 | 0.075 | -7.525 | 4.203 |
| | | Men | m1 | 0.621 | 2.427 | 4.890 |
| | | | m2 | 0.567 | 2.802 | 4.875 |
| | | | m3 | 0.300 | 5.952 | 5.710 |
| | | | m4 | 0.393 | 4.891 | 5.695 |
| | | Women | m1 | 0.000 | -19.177 | 5.340 |
| | | | m2 | 0.001 | -18.458 | 5.423 |
| | | | m3 | 0.003 | -17.506 | 5.846 |
| | | | m4 | 0.003 | -18.318 | 5.975 |
| Nighttime SBP CoV | *Bacteroides plebeius* | All | m1 | 0.031 | -4.685 | 2.165 |
| | | | m2 | 0.057 | -4.164 | 2.175 |
| | | | m3 | 0.065 | -4.538 | 2.445 |
| | | | m4 | 0.050 | -4.865 | 2.470 |
| | | Men | m1 | 0.803 | -0.677 | 2.700 |
| | | | m2 | 0.767 | -0.811 | 2.723 |
| | | | m3 | 0.975 | -0.093 | 3.031 |
| | | | m4 | 0.877 | 0.489 | 3.159 |
| | | Women | m1 | 0.007 | -9.516 | 3.484 |
| | | | m2 | 0.019 | -8.010 | 3.378 |
| | | | m3 | 0.016 | -9.522 | 3.870 |
| | | | m4 | 0.035 | -8.441 | 3.954 |
| | *Blautia hansenii* | All | m1 | 0.428 | 4.796 | 6.034 |
| | | | m2 | 0.329 | 5.974 | 6.111 |
| | | | m3 | 0.063 | 12.686 | 6.794 |
| | | | m4 | 0.041 | 14.892 | 7.234 |
| | | Men | m1 | 0.113 | 9.610 | 6.010 |
| | | | m2 | 0.134 | 9.270 | 6.143 |
| | | | m3 | 0.009 | 16.694 | 6.270 |
| | | | m4 | 0.018 | 15.947 | 6.599 |
| | | Women | m1 | 0.332 | -17.637 | 18.129 |
| | | | m2 | 0.078 | -30.840 | 17.381 |
| | | | m3 | 0.265 | -23.072 | 20.569 |
| | | | m4 | 0.182 | -38.633 | 28.737 |
| | *Blautia wexlerae* | All | m1 | 0.086 | -6.727 | 3.896 |
| | | | m2 | 0.096 | -6.436 | 3.850 |
| | | | m3 | 0.043 | -8.153 | 4.004 |
| | | | m4 | 0.023 | -9.381 | 4.104 |
| | | Men | m1 | 0.857 | 1.148 | 6.345 |
| | | | m2 | 0.881 | 0.966 | 6.458 |
| | | | m3 | 0.977 | -0.206 | 7.091 |
| | | | m4 | 0.814 | -1.707 | 7.244 |

| | | | | | | |
|---|---|---|---|---|---|---|
| | | Women | m1 | 0.036 | -10.652 | 5.029 |
| | | | m2 | 0.023 | -10.995 | 4.768 |
| | | | m3 | 0.022 | -12.202 | 5.252 |
| | | | m4 | 0.008 | -14.610 | 5.364 |
| | *Coprococcus catus* | All | m1 | 0.224 | -7.697 | 6.313 |
| | | | m2 | 0.219 | -7.703 | 6.254 |
| | | | m3 | 0.014 | -16.856 | 6.820 |
| | | | m4 | 0.005 | -19.951 | 6.998 |
| | | Men | m1 | 0.330 | 8.236 | 8.424 |
| | | | m2 | 0.352 | 7.959 | 8.521 |
| | | | m3 | 0.734 | 3.076 | 9.007 |
| | | | m4 | 0.949 | -0.638 | 9.874 |
| | | Women | m1 | 0.014 | -22.821 | 9.187 |
| | | | m2 | 0.043 | -18.197 | 8.909 |
| | | | m3 | 0.012 | -26.314 | 10.296 |
| | | | m4 | 0.013 | -26.665 | 10.507 |
| | *Escherichia coli* | All | m1 | 0.057 | 5.464 | 2.857 |
| | | | m2 | 0.086 | 4.902 | 2.843 |
| | | | m3 | 0.288 | 3.683 | 3.460 |
| | | | m4 | 0.378 | 3.112 | 3.522 |
| | | Men | m1 | 0.007 | 14.228 | 5.202 |
| | | | m2 | 0.007 | 14.295 | 5.231 |
| | | | m3 | 0.883 | 1.066 | 7.247 |
| | | | m4 | 0.665 | 3.256 | 7.495 |
| | | Women | m1 | 0.497 | 2.434 | 3.574 |
| | | | m2 | 0.701 | 1.315 | 3.420 |
| | | | m3 | 0.213 | 5.431 | 4.333 |
| | | | m4 | 0.290 | 4.668 | 4.382 |
| | *Megamonas funiformis* | All | m1 | 0.541 | 2.154 | 3.519 |
| | | | m2 | 0.616 | 1.763 | 3.508 |
| | | | m3 | 0.039 | 9.062 | 4.358 |
| | | | m4 | 0.051 | 8.690 | 4.429 |
| | | Men | m1 | 0.720 | -1.586 | 4.407 |
| | | | m2 | 0.688 | -1.791 | 4.445 |
| | | | m3 | 0.445 | 3.561 | 4.644 |
| | | | m4 | 0.545 | 2.885 | 4.742 |
| | | Women | m1 | 0.189 | 7.404 | 5.612 |
| | | | m2 | 0.272 | 5.920 | 5.364 |
| | | | m3 | 0.007 | 22.245 | 8.125 |
| | | | m4 | 0.017 | 20.741 | 8.502 |
| | *Megamonas hypermegale* | All | m1 | 0.743 | 1.910 | 5.829 |
| | | | m2 | 0.642 | 2.745 | 5.898 |
| | | | m3 | 0.206 | 8.427 | 6.647 |

| | | | | | | |
|---|---|---|---|---|---|---|
| | | | m4 | 0.264 | 7.669 | 6.846 |
| | | Men | m1 | 0.425 | -5.325 | 6.644 |
| | | | m2 | 0.362 | -6.195 | 6.771 |
| | | | m3 | 0.376 | -6.542 | 7.344 |
| | | | m4 | 0.509 | -4.978 | 7.499 |
| | | Women | m1 | 0.081 | 19.275 | 10.968 |
| | | | m2 | 0.127 | 16.144 | 10.514 |
| | | | m3 | 0.006 | 35.392 | 12.640 |
| | | | m4 | 0.016 | 32.904 | 13.345 |
| | *Roseburia intestinalis* | All | m1 | 0.035 | 7.412 | 3.497 |
| | | | m2 | 0.063 | 6.492 | 3.476 |
| | | | m3 | 0.220 | 4.757 | 3.864 |
| | | | m4 | 0.201 | 5.082 | 3.962 |
| | | Men | m1 | 0.003 | 13.657 | 4.521 |
| | | | m2 | 0.004 | 13.468 | 4.563 |
| | | | m3 | 0.023 | 11.133 | 4.810 |
| | | | m4 | 0.032 | 10.973 | 5.015 |
| | | Women | m1 | 0.698 | 2.033 | 5.221 |
| | | | m2 | 0.842 | -1.007 | 5.035 |
| | | | m3 | 0.807 | -1.416 | 5.783 |
| | | | m4 | 0.735 | -2.034 | 5.981 |
| | *Ruminococcus bicirculans* | All | m1 | 0.607 | 1.558 | 3.023 |
| | | | m2 | 0.542 | 1.844 | 3.021 |
| | | | m3 | 0.845 | 0.626 | 3.193 |
| | | | m4 | 0.784 | 0.885 | 3.221 |
| | | Men | m1 | 0.007 | 11.805 | 4.322 |
| | | | m2 | 0.004 | 13.196 | 4.421 |
| | | | m3 | 0.006 | 12.656 | 4.490 |
| | | | m4 | 0.007 | 12.529 | 4.487 |
| | | Women | m1 | 0.193 | -5.469 | 4.179 |
| | | | m2 | 0.280 | -4.338 | 3.995 |
| | | | m3 | 0.397 | -3.774 | 4.435 |
| | | | m4 | 0.521 | -2.890 | 4.483 |
| Nighttime DBP CoV | *Adlercreutzia equolifaciens* | All | m1 | 0.017 | -21.801 | 9.085 |
| | | | m2 | 0.021 | -20.987 | 9.064 |
| | | | m3 | 0.030 | -21.669 | 9.935 |
| | | | m4 | 0.008 | -26.958 | 10.002 |
| | | Men | m1 | 0.139 | -22.888 | 15.354 |
| | | | m2 | 0.143 | -22.671 | 15.376 |
| | | | m3 | 0.413 | -14.659 | 17.822 |
| | | | m4 | 0.458 | -13.441 | 18.012 |
| | | Women | m1 | 0.064 | -20.838 | 11.154 |
| | | | m2 | 0.111 | -17.514 | 10.924 |

| | | | | | | |
|---|---|---|---|---|---|---|
| | | | m3 | 0.046 | -24.622 | 12.178 |
| | | | m4 | 0.020 | -29.927 | 12.615 |
| | *Asaccharobacter celatus* | All | m1 | 0.036 | -31.658 | 15.044 |
| | | | m2 | 0.045 | -30.248 | 15.010 |
| | | | m3 | 0.040 | -37.354 | 18.037 |
| | | | m4 | 0.009 | -47.863 | 18.218 |
| | | Men | m1 | 0.088 | -41.787 | 24.253 |
| | | | m2 | 0.100 | -40.387 | 24.314 |
| | | | m3 | 0.352 | -29.620 | 31.669 |
| | | | m4 | 0.357 | -29.630 | 31.992 |
| | | Women | m1 | 0.212 | -23.919 | 19.072 |
| | | | m2 | 0.333 | -18.145 | 18.662 |
| | | | m3 | 0.064 | -41.992 | 22.380 |
| | | | m4 | 0.028 | -51.969 | 23.254 |
| | *Blautia hansenii* | All | m1 | 0.047 | 14.914 | 7.467 |
| | | | m2 | 0.087 | 13.183 | 7.664 |
| | | | m3 | 0.021 | 19.626 | 8.436 |
| | | | m4 | 0.025 | 20.231 | 8.945 |
| | | Men | m1 | 0.011 | 21.681 | 8.371 |
| | | | m2 | 0.019 | 20.263 | 8.535 |
| | | | m3 | 0.002 | 28.744 | 8.862 |
| | | | m4 | 0.003 | 28.135 | 9.240 |
| | | Women | m1 | 0.084 | -35.073 | 20.161 |
| | | | m2 | 0.019 | -46.851 | 19.694 |
| | | | m3 | 0.056 | -42.853 | 22.195 |
| | | | m4 | 0.023 | -72.678 | 31.417 |
| | *Fusicatenibacter saccharivorans* | All | m1 | 0.244 | -3.402 | 2.916 |
| | | | m2 | 0.222 | -3.569 | 2.915 |
| | | | m3 | 0.095 | -5.139 | 3.067 |
| | | | m4 | 0.141 | -4.568 | 3.089 |
| | | Men | m1 | 0.646 | 1.971 | 4.284 |
| | | | m2 | 0.790 | 1.156 | 4.335 |
| | | | m3 | 0.985 | -0.081 | 4.446 |
| | | | m4 | 0.943 | 0.317 | 4.446 |
| | | Women | m1 | 0.032 | -8.588 | 3.962 |
| | | | m2 | 0.006 | -10.658 | 3.847 |
| | | | m3 | 0.002 | -13.344 | 4.137 |
| | | | m4 | 0.004 | -12.748 | 4.319 |
| | *Megamonas hypermegale* | All | m1 | 0.096 | 12.080 | 7.223 |
| | | | m2 | 0.171 | 10.160 | 7.401 |
| | | | m3 | 0.027 | 18.358 | 8.223 |
| | | | m4 | 0.006 | 23.009 | 8.341 |
| | | Men | m1 | 0.174 | 12.835 | 9.371 |

| | | | | | | |
|---|---|---|---|---|---|---|
| | | | m2 | 0.248 | 11.059 | 9.526 |
| | | | m3 | 0.202 | 13.540 | 10.527 |
| | | | m4 | 0.103 | 17.458 | 10.570 |
| | | Women | m1 | 0.523 | 7.952 | 12.427 |
| | | | m2 | 0.721 | 4.339 | 12.138 |
| | | | m3 | 0.084 | 24.678 | 14.136 |
| | | | m4 | 0.036 | 31.939 | 14.972 |

Legend. Gut bacterial species associated with BP variability indices determined by linear regression analysis performed systematically for each bacterial species under sex-stratified cohorts and four models for covariate adjustment, m1: no covariate adjustment, m2: age; sex; BMI, m3: m2 + sodium intake based on spot urine analysis; serum glucose, triglyceride, HDL, and LDL cholesterol; smoking status; menopause status; liver steatosis by CAP score, m4: m3 + 24-hour mean SBP or DBP; sleep latency. Abbreviations: BP, blood pressure; CoV, coefficient of variation; DBP, diastolic blood pressure; HDL, high-density lipoprotein; LDL, low-density lipoprotein; SBP, systolic blood pressure.

**Table S5. Association between 24-hour BP variability indices and plasma SCFAs**

| 24-hour BP Variability Index | Plasma SCFA | Sex | Model (m) | *P*-value | Coefficient | Standard error |
|---|---|---|---|---|---|---|
| 24-hour SBP CoV | Iso-butyric Acid | All | m1 | 0.006 | -0.066 | 0.024 |
| | | | m2 | 0.019 | -0.056 | 0.024 |
| | | | m3 | 0.004 | -0.075 | 0.026 |
| | | | m4 | 0.011 | -0.068 | 0.027 |
| | | Men | m1 | 0.188 | -0.031 | 0.023 |
| | | | m2 | 0.204 | -0.030 | 0.023 |
| | | | m3 | 0.076 | -0.041 | 0.023 |
| | | | m4 | 0.075 | -0.040 | 0.022 |
| | | Women | m1 | 0.038 | -0.089 | 0.042 |
| | | | m2 | 0.028 | -0.094 | 0.042 |
| | | | m3 | 0.013 | -0.126 | 0.050 |
| | | | m4 | 0.021 | -0.121 | 0.052 |
| Daytime SBP CoV | Iso-butyric Acid | All | m1 | 0.019 | -0.055 | 0.023 |
| | | | m2 | 0.065 | -0.043 | 0.023 |
| | | | m3 | 0.011 | -0.064 | 0.025 |
| | | | m4 | 0.013 | -0.065 | 0.026 |
| | | Men | m1 | 0.313 | -0.026 | 0.026 |
| | | | m2 | 0.353 | -0.024 | 0.026 |
| | | | m3 | 0.126 | -0.040 | 0.026 |
| | | | m4 | 0.089 | -0.044 | 0.025 |
| | | Women | m1 | 0.080 | -0.069 | 0.039 |
| | | | m2 | 0.087 | -0.068 | 0.039 |
| | | | m3 | 0.019 | -0.111 | 0.046 |
| | | | m4 | 0.022 | -0.113 | 0.049 |
| Nighttime SBP CoV | Butyric Acid | All | m1 | 0.289 | -0.075 | 0.071 |
| | | | m2 | 0.287 | -0.075 | 0.070 |
| | | | m3 | 0.451 | -0.057 | 0.076 |
| | | | m4 | 0.318 | -0.077 | 0.077 |
| | | Men | m1 | 0.887 | 0.012 | 0.083 |
| | | | m2 | 0.945 | 0.006 | 0.085 |
| | | | m3 | 0.831 | -0.018 | 0.085 |
| | | | m4 | 0.960 | -0.004 | 0.088 |
| | | Women | m1 | 0.110 | -0.193 | 0.120 |
| | | | m2 | 0.039 | -0.236 | 0.113 |
| | | | m3 | 0.326 | -0.133 | 0.135 |
| | | | m4 | 0.254 | -0.157 | 0.136 |
| 24-hour DBP CoV | Propionic Acid | All | m1 | 0.121 | -0.003 | 0.002 |
| | | | m2 | 0.420 | -0.002 | 0.002 |
| | | | m3 | 0.728 | -0.001 | 0.002 |
| | | | m4 | 0.801 | 0.001 | 0.002 |

| | | | | | | |
|---|---|---|---|---|---|---|
| | | Men | m1 | 0.532 | 0.001 | 0.002 |
| | | | m2 | 0.590 | 0.001 | 0.002 |
| | | | m3 | 0.533 | 0.002 | 0.003 |
| | | | m4 | 0.557 | 0.001 | 0.002 |
| | | Women | m1 | 0.037 | -0.007 | 0.003 |
| | | | m2 | 0.050 | -0.007 | 0.003 |
| | | | m3 | 0.139 | -0.007 | 0.005 |
| | | | m4 | 0.300 | -0.005 | 0.005 |
| Nighttime DBP CoV | Butyric Acid | All | m1 | 0.149 | -0.128 | 0.088 |
| | | | m2 | 0.096 | -0.148 | 0.088 |
| | | | m3 | 0.137 | -0.140 | 0.094 |
| | | | m4 | 0.176 | -0.130 | 0.096 |
| | | Men | m1 | 0.720 | -0.043 | 0.120 |
| | | | m2 | 0.604 | -0.063 | 0.122 |
| | | | m3 | 0.560 | -0.074 | 0.126 |
| | | | m4 | 0.653 | -0.059 | 0.130 |
| | | Women | m1 | 0.045 | -0.273 | 0.135 |
| | | | m2 | 0.018 | -0.311 | 0.129 |
| | | | m3 | 0.214 | -0.183 | 0.146 |
| | | | m4 | 0.263 | -0.170 | 0.151 |
| Nondippers vs dippers | Butyric Acid | All | m1 | 0.297 | 0.046 | 0.044 |
| | | | m2 | 0.363 | 0.042 | 0.046 |
| | | | m3 | 0.354 | 0.048 | 0.052 |
| | | | m4 | 0.252 | 0.063 | 0.055 |
| | | Men | m1 | 0.023 | 0.174 | 0.077 |
| | | | m2 | 0.028 | 0.173 | 0.079 |
| | | | m3 | 0.017 | 0.327 | 0.137 |
| | | | m4 | 0.009 | 0.503 | 0.192 |
| | | Women | m1 | 0.199 | -0.085 | 0.066 |
| | | | m2 | 0.219 | -0.082 | 0.066 |
| | | | m3 | 0.099 | -0.135 | 0.082 |
| | | | m4 | 0.135 | -0.127 | 0.085 |

Legend. Plasma SCFAs associations with BP variability indices determined by linear regression analysis performed systematically for each bacterial species under sex-stratified cohorts, and four models for covariate adjustment, m1: no covariate adjustment, m2: age; sex; BMI, m3: m2 + sodium intake based on spot urine analysis; serum glucose, triglyceride, HDL, and LDL cholesterol; smoking status; menopause status; liver steatosis by CAP score, m4: m3 + 24-hour mean SBP or DBP; sleep latency. Abbreviations: BP, blood pressure; CoV, coefficient of variation; DBP, diastolic blood pressure; HDL, high-density lipoprotein; LDL, low-density lipoprotein; SBP, systolic blood pressure; SCFA, short-chain fatty acid.

**Table S6. Association between plasma SCFAs and GM species**

| Plasma SCFA | GM species | Sex | Model (m) | *P*-value | Coefficient | Standard error |
|---|---|---|---|---|---|---|
| Iso-butyric Acid | *Bacteroides dorei* | All | m1 | 0.688 | 1.824 | 4.535 |
| | | | m2 | 0.426 | 3.651 | 4.575 |
| | | | m3 | 0.455 | 3.616 | 4.827 |
| | | | m4 | 0.629 | 2.378 | 4.921 |
| | | Men | m1 | 0.193 | -10.536 | 8.049 |
| | | | m2 | 0.212 | -10.191 | 8.118 |
| | | | m3 | 0.110 | -14.597 | 9.041 |
| | | | m4 | 0.083 | -15.986 | 9.100 |
| | | Women | m1 | 0.021 | 11.976 | 5.126 |
| | | | m2 | 0.019 | 12.254 | 5.169 |
| | | | m3 | 0.003 | 16.145 | 5.324 |
| | | | m4 | 0.011 | 14.374 | 5.543 |
| | *Bacteroides coprocola* | All | m1 | 0.108 | 15.681 | 9.719 |
| | | | m2 | 0.150 | 14.113 | 9.764 |
| | | | m3 | 0.035 | 22.822 | 10.737 |
| | | | m4 | 0.027 | 23.984 | 10.785 |
| | | Men | m1 | 0.040 | 32.268 | 15.517 |
| | | | m2 | 0.039 | 32.716 | 15.646 |
| | | | m3 | 0.004 | 55.629 | 18.695 |
| | | | m4 | 0.003 | 58.530 | 18.859 |
| | | Women | m1 | 0.885 | -1.729 | 11.979 |
| | | | m2 | 0.932 | 1.052 | 12.378 |
| | | | m3 | 0.785 | 3.658 | 13.361 |
| | | | m4 | 0.671 | 5.689 | 13.328 |
| | *Faecalibacterium prausnitzii* | All | m1 | 0.199 | 4.845 | 3.763 |
| | | | m2 | 0.145 | 5.484 | 3.751 |
| | | | m3 | 0.224 | 5.125 | 4.197 |
| | | | m4 | 0.150 | 6.122 | 4.230 |
| | | Men | m1 | 0.099 | 9.737 | 5.858 |
| | | | m2 | 0.079 | 10.546 | 5.948 |
| | | | m3 | 0.064 | 13.451 | 7.159 |
| | | | m4 | 0.042 | 15.273 | 7.396 |
| | | Women | m1 | 0.849 | 0.904 | 4.730 |
| | | | m2 | 0.794 | 1.244 | 4.754 |
| | | | m3 | 0.419 | -4.359 | 5.366 |
| | | | m4 | 0.575 | -3.061 | 5.447 |
| Butyric Acid | *Bacteroides dorei* | All | m1 | 0.521 | -1.671 | 2.600 |

| | | | | | | |
|---|---|---|---|---|---|---|
| | | | m2 | 0.735 | -0.899 | 2.651 |
| | | | m3 | 0.538 | -1.792 | 2.905 |
| | | | m4 | 0.489 | -2.079 | 2.999 |
| | | Men | m1 | 0.154 | -7.621 | 5.303 |
| | | | m2 | 0.167 | -7.404 | 5.312 |
| | | | m3 | 0.036 | -12.292 | 5.724 |
| | | | m4 | 0.048 | -11.748 | 5.809 |
| | | Women | m1 | 0.470 | 1.970 | 2.717 |
| | | | m2 | 0.498 | 1.886 | 2.772 |
| | | | m3 | 0.398 | 2.634 | 3.102 |
| | | | m4 | 0.484 | 2.319 | 3.298 |
| | *Bacteroides coprocola* | All | m1 | 0.168 | 7.885 | 5.704 |
| | | | m2 | 0.219 | 7.199 | 5.842 |
| | | | m3 | 0.062 | 12.716 | 6.758 |
| | | | m4 | 0.070 | 12.518 | 6.853 |
| | | Men | m1 | 0.091 | 17.794 | 10.427 |
| | | | m2 | 0.135 | 15.998 | 10.603 |
| | | | m3 | 0.008 | 35.939 | 13.056 |
| | | | m4 | 0.012 | 34.704 | 13.371 |
| | | Women | m1 | 0.889 | 0.880 | 6.261 |
| | | | m2 | 0.684 | 2.686 | 6.579 |
| | | | m3 | 0.979 | -0.203 | 7.593 |
| | | | m4 | 0.994 | -0.061 | 7.797 |
| | *Firmicutes bacterium CAG 83* | All | m1 | 0.174 | 6.956 | 5.099 |
| | | | m2 | 0.081 | 9.309 | 5.301 |
| | | | m3 | 0.021 | 13.601 | 5.831 |
| | | | m4 | 0.017 | 14.363 | 5.964 |
| | | Men | m1 | 0.933 | -0.928 | 11.074 |
| | | | m2 | 0.887 | -1.591 | 11.129 |
| | | | m3 | 0.499 | 9.249 | 13.591 |
| | | | m4 | 0.496 | 9.387 | 13.713 |
| | | Women | m1 | 0.010 | 13.640 | 5.226 |
| | | | m2 | 0.005 | 15.134 | 5.322 |
| | | | m3 | 0.008 | 15.868 | 5.841 |
| | | | m4 | 0.008 | 16.394 | 6.032 |
| Propionic Acid | *Bacteroides dorei* | All | m1 | 0.667 | -57.291 | 132.772 |
| | | | m2 | 0.624 | 63.981 | 130.449 |
| | | | m3 | 0.614 | 68.434 | 135.383 |
| | | | m4 | 0.694 | 55.018 | 139.558 |
| | | Men | m1 | 0.173 | -354.436 | 257.536 |
| | | | m2 | 0.175 | -350.651 | 256.143 |
| | | | m3 | 0.427 | -239.646 | 299.443 |
| | | | m4 | 0.474 | -222.149 | 308.024 |

| | | | | | | |
|---|---|---|---|---|---|---|
| | | Women | m1 | 0.050 | 262.036 | 131.476 |
| | | | m2 | 0.057 | 256.838 | 132.981 |
| | | | m3 | 0.091 | 205.906 | 119.863 |
| | | | m4 | 0.061 | 235.372 | 123.267 |
| | *Bacteroides plebeius* | All | m1 | 0.048 | 225.223 | 113.103 |
| | | | m2 | 0.142 | 162.908 | 110.419 |
| | | | m3 | 0.379 | 115.906 | 131.155 |
| | | | m4 | 0.386 | 115.946 | 133.161 |
| | | Men | m1 | 0.085 | 298.759 | 171.492 |
| | | | m2 | 0.094 | 290.961 | 171.487 |
| | | | m3 | 0.098 | 446.120 | 264.544 |
| | | | m4 | 0.101 | 457.710 | 273.457 |
| | | Women | m1 | 0.848 | 25.971 | 135.268 |
| | | | m2 | 0.799 | 35.081 | 137.586 |
| | | | m3 | 0.226 | -160.935 | 131.628 |
| | | | m4 | 0.149 | -191.152 | 130.529 |
| | *Faecalibacterium prausnitzii* | All | m1 | 0.395 | -91.550 | 107.262 |
| | | | m2 | 0.685 | -41.990 | 103.438 |
| | | | m3 | 0.811 | -27.459 | 114.870 |
| | | | m4 | 0.977 | -3.376 | 117.895 |
| | | Men | m1 | 0.576 | 91.201 | 162.361 |
| | | | m2 | 0.367 | 148.996 | 164.133 |
| | | | m3 | 0.281 | 227.202 | 208.278 |
| | | | m4 | 0.330 | 217.599 | 221.106 |
| | | Women | m1 | 0.097 | -209.658 | 124.891 |
| | | | m2 | 0.102 | -209.089 | 126.524 |
| | | | m3 | 0.008 | -337.849 | 122.509 |
| | | | m4 | 0.014 | -317.859 | 124.879 |
| Acetic Acid | *Bacteroides plebeius* | All | m1 | 0.092 | -83.892 | 49.594 |
| | | | m2 | 0.027 | -109.334 | 49.157 |
| | | | m3 | 0.029 | -125.243 | 56.934 |
| | | | m4 | 0.025 | -131.267 | 58.047 |
| | | Men | m1 | 0.332 | -74.606 | 76.498 |
| | | | m2 | 0.240 | -88.675 | 75.136 |
| | | | m3 | 0.291 | -106.680 | 100.321 |
| | | | m4 | 0.195 | -137.120 | 104.825 |
| | | Women | m1 | 0.109 | -105.235 | 65.277 |
| | | | m2 | 0.040 | -132.935 | 64.013 |
| | | | m3 | 0.043 | -145.992 | 71.255 |
| | | | m4 | 0.038 | -154.627 | 73.500 |
| | *Ruthenibacterium lactatiformans* | All | m1 | 0.068 | 250.039 | 136.336 |
| | | | m2 | 0.036 | 289.076 | 137.415 |
| | | | m3 | 0.020 | 351.589 | 149.354 |

| | | | | | | |
|---|---|---|---|---|---|---|
| | | | m4 | 0.023 | 353.191 | 153.814 |
| | | Men | m1 | 0.145 | 452.795 | 308.849 |
| | | | m2 | 0.175 | 416.371 | 304.656 |
| | | | m3 | 0.157 | 509.826 | 357.133 |
| | | | m4 | 0.148 | 529.880 | 362.563 |
| | | Women | m1 | 0.152 | 201.497 | 139.729 |
| | | | m2 | 0.062 | 261.798 | 139.155 |
| | | | m3 | 0.008 | 428.774 | 159.516 |
| | | | m4 | 0.010 | 440.412 | 168.348 |
| Total SCFAs | *Bacteroides dorei* | All | m1 | 0.956 | -9.265 | 166.299 |
| | | | m2 | 0.366 | 146.514 | 161.721 |
| | | | m3 | 0.395 | 142.867 | 167.200 |
| | | | m4 | 0.515 | 113.276 | 173.229 |
| | | Men | m1 | 0.171 | -436.343 | 315.524 |
| | | | m2 | 0.183 | -422.998 | 314.547 |
| | | | m3 | 0.463 | -275.704 | 372.641 |
| | | | m4 | 0.517 | -252.044 | 385.608 |
| | | Women | m1 | 0.026 | 385.759 | 170.351 |
| | | | m2 | 0.025 | 389.597 | 170.735 |
| | | | m3 | 0.057 | 304.054 | 156.419 |
| | | | m4 | 0.052 | 321.294 | 161.499 |
| | *Faecalibacterium prausnitzii* | All | m1 | 0.585 | -73.920 | 134.991 |
| | | | m2 | 0.918 | -13.363 | 129.306 |
| | | | m3 | 0.775 | -40.829 | 142.258 |
| | | | m4 | 0.940 | -11.040 | 147.077 |
| | | Men | m1 | 0.304 | 202.220 | 195.512 |
| | | | m2 | 0.202 | 253.588 | 197.098 |
| | | | m3 | 0.165 | 358.447 | 253.839 |
| | | | m4 | 0.212 | 351.866 | 277.273 |
| | | Women | m1 | 0.132 | -253.725 | 166.617 |
| | | | m2 | 0.100 | -280.528 | 168.418 |
| | | | m3 | 0.001 | -565.055 | 154.571 |
| | | | m4 | 0.001 | -546.890 | 158.464 |
| | *Ruthenibacterium lactatiformans* | All | m1 | 0.454 | -266.306 | 354.447 |
| | | | m2 | 0.781 | 97.312 | 349.213 |
| | | | m3 | 0.447 | 284.772 | 373.116 |
| | | | m4 | 0.551 | 230.082 | 384.575 |
| | | Men | m1 | 0.659 | 339.409 | 766.371 |
| | | | m2 | 0.687 | 310.510 | 767.329 |
| | | | m3 | 0.736 | 322.298 | 950.002 |
| | | | m4 | 0.640 | 471.911 | 1000.187 |
| | | Women | m1 | 0.830 | -77.428 | 359.872 |
| | | | m2 | 0.932 | 31.412 | 365.694 |

| | | | | | | |
|---|---|---|---|---|---|---|
| | | | m3 | 0.012 | 964.322 | 372.234 |
| | | | m4 | 0.029 | 873.153 | 387.126 |

Legend. Linear regression analysis of associations between plasma SCFAs and gut bacterial species with sex-stratification under covariate-adjusted models (m), m1: no covariate adjustment, m2: age; sex; BMI, m3: m2 + sodium intake based on spot urine analysis; serum glucose, triglyceride, HDL, and LDL cholesterol; smoking status; menopause status; liver steatosis by CAP score, m4: m3 + 24-hour mean SBP or DBP; sleep latency. Abbreviations: HDL, high-density lipoprotein; LDL, low-density lipoprotein; SCFA, short-chain fatty acid.

##### Table S7. Association between GM and nighttime dipping status

| Nighttime dipping status | GM species | Sex | Model (m) | *P*-value | Coefficient | Standard error |
|---|---|---|---|---|---|---|
| Nondippers vs. Dippers | *Bacteroides intestinalis* | All | m1 | 0.050 | -9.540 | 4.871 |
| | | | m2 | 0.047 | -10.035 | 5.049 |
| | | | m3 | 0.026 | -13.442 | 6.035 |
| | | | m4 | 0.036 | -13.313 | 6.352 |
| | | Men | m1 | 0.312 | 12.295 | 12.173 |
| | | | m2 | 0.340 | 11.639 | 12.206 |
| | | | m3 | 0.485 | 11.061 | 15.836 |
| | | | m4 | 0.520 | 10.609 | 16.487 |
| | | Women | m1 | 0.012 | -16.620 | 6.598 |
| | | | m2 | 0.006 | -18.214 | 6.671 |
| | | | m3 | 0.005 | -22.168 | 7.948 |
| | | | m4 | 0.006 | -22.137 | 8.130 |
| | *Alistipes finegoldii* | All | m1 | 0.247 | -3.876 | 3.345 |
| | | | m2 | 0.174 | -4.669 | 3.435 |
| | | | m3 | 0.110 | -6.173 | 3.867 |
| | | | m4 | 0.200 | -5.084 | 3.971 |
| | | Men | m1 | 0.504 | 4.188 | 6.266 |
| | | | m2 | 0.485 | 4.479 | 6.412 |
| | | | m3 | 0.628 | 3.753 | 7.743 |
| | | | m4 | 0.569 | 4.468 | 7.844 |
| | | Women | m1 | 0.069 | -8.530 | 4.690 |
| | | | m2 | 0.040 | -9.958 | 4.855 |
| | | | m3 | 0.005 | -18.902 | 6.660 |
| | | | m4 | 0.007 | -17.831 | 6.557 |
| | *Eubacterium rectale* | All | m1 | 0.324 | 0.928 | 0.940 |
| | | | m2 | 0.201 | 1.223 | 0.956 |
| | | | m3 | 0.094 | 1.848 | 1.102 |
| | | | m4 | 0.093 | 1.922 | 1.146 |
| | | Men | m1 | 0.065 | 3.345 | 1.815 |
| | | | m2 | 0.050 | 3.681 | 1.875 |
| | | | m3 | 0.016 | 5.747 | 2.391 |
| | | | m4 | 0.009 | 6.905 | 2.631 |
| | | Women | m1 | 0.899 | 0.141 | 1.109 |
| | | | m2 | 0.836 | 0.233 | 1.126 |
| | | | m3 | 0.664 | 0.601 | 1.384 |
| | | | m4 | 0.746 | 0.465 | 1.437 |

Legend. Logistic regression model results displaying the associations of GM with non-dippers vs. dippers. Sex-stratified analysis was performed systematically for each bacterial species under four models (m) of covariate adjustment, m1: no covariate adjustment, m2: age; sex; BMI, m3: m2 + sodium intake based on spot urine analysis; serum glucose, triglyceride, HDL, and LDL cholesterol; smoking status; menopause status; liver steatosis by CAP score, m4: m3 + 24-hour mean SBP or DBP; sleep latency. Only bacterial species with at least one statistically significant ($p <= 0.01$) association under any model or sex-stratified analysis were included in this table. Abbreviations: HDL, high-density lipoprotein; LDL, low-density lipoprotein.
